# A multivariate outcome test of covariance

**DOI:** 10.1101/2023.09.20.558234

**Authors:** Christophe Boetto, Arthur Frouin, Léo Henches, Antoine Auvergne, Yuka Suzuki, Etienne Patin, Marius Bredon, Alec Chiu, Milieu Interieur Consortium, Sriram Sankararaman, Noah Zaitlen, Sean P. Kennedy, Lluis Quintana-Murci, Darragh Duffy, Harry Sokol, Hugues Aschard

## Abstract

Multivariate analysis is becoming central in studies investigating high-throughput molecular data, yet, some important features of these data are seldom explored. Here, we present MANOCCA (Multivariate Analysis of Conditional CovAriance), a powerful method to test for the effect of a predictor on the covariance matrix of a multivariate outcome. The proposed test is by construction orthogonal to tests based on the mean and variance, and is able to capture effects that are missed by both approaches. We first compare the performances of MANOCCA with existing correlation-based methods and show that MANOCCA is the only test correctly calibrated in simulation mimicking omics data. We then investigate the impact of reducing the dimensionality of the data using principal component analysis when the sample size is smaller than the number of pairwise covariance terms analysed. We show that, in many realistic scenarios, the maximum power can be achieved with a limited number of components. Finally, we apply MANOCCA to 1,000 healthy individuals from the Milieu Interieur cohort, to assess the effect of health, lifestyle and genetic factors on the covariance of two sets of phenotypes, blood biomarkers and flow cytometry-based immune phenotypes. Our analyses identify significant associations between multiple factors and the covariance of both omics data.

## Introduction

Human cohorts commonly collect high-dimensional phenotypic data, including high-throughput omics, extended medical information, and biomarkers^1,2^. A variety of multivariate approaches have been developed to leverage this wealth of data^3-6^. The joint analysis of multiple outcomes can increase statistical power to detect associations^7,8^, help deciphering complex biological processes through clustering approaches^9^, or improve the prediction accuracy of an outcome of interest^10^. Regarding association testing, existing methods and application have mostly focused on testing the impact of predictors of interest on the mean of a multivariate outcome, typically using a composite null hypothesis such as implemented in a multivariate ANOVA. Conversely, methods to investigate other components of multivariate outcomes remain sparse. One of such components of multivariate outcomes is correlation, which is commonly present in omics data. Although methods to investigate predictors associated with the correlation between multiple outcomes exist^11-14^, their performance and robustness have not been assessed, and their efficiency in large-scale agnostic screenings remains unknown. Moreover, they carry substantial inherent limitations, including restriction to binary factors and no adjustment for covariates.

Here, we present a new approach, named MANOCCA (Multivariate ANalysis Of Conditional Covariance), that enables the identification of both categorical and continuous predictors associated with changes in the covariance matrix of a multivariate outcome while allowing for covariates adjustment. We first introduce the key principles and the main characteristics of the approach, and demonstrate that, in most realistic scenarios, MANOCCA can outperform existing approaches showing stronger power and robustness. We then describe the challenges faced when analysing high-dimensional data, and present a robust solution based on principal components analysis. We next investigate the power of MANOCCA conditional on alternative parametrizations, providing guidelines for real data application across various settings. Finally, we illustrate the method by studying health, lifestyle and genetic factors associated with variability of blood biomarkers and flow cytometry-based immune phenotypes using data from 1,000 healthy subjects from the *Milieu Intérieur* cohort.

## Methods

### The MANOCCA approach

Previous work^15^ showed that variability in the correlation between two standardized outcomes *Y*_1_ and *Y*_2_ can be investigated through the element-wise product of those outcomes. The Pearson correlation coefficient between *Y*_1_ and *Y*_2_ is expressed as 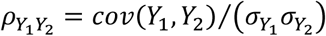, with *cov*(*Y*_1_, *Y*_2_) = 𝔼[*Y*_1_*Y*_2_] − 𝔼[*Y*_1_]𝔼[*Y*_2_]. For standardized outcomes and a sample size *N*, it can be re-expressed as the average of the element-wise product across individuals: 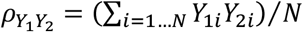. It follows that the effect of a predictor *X* on *cor*(*Y*_1_, *Y*_2_) can be tested using a standard least-squares regression framework where *X* is treated as a predictor and the product *Y*_1_*Y*_2_ as the outcome. One can easily demonstrate that, under reasonable assumptions, this test is independent of mean and variance effect. Consider the following models: *Y*_1_ = *α*_1_*U* + *β*_1_*X* + *ϵ*_1_ and *Y*_2_ = *α*_2_*U* + *β*_2_*X* + *ϵ*_2_, where *Y*_1_ and *Y*_2_ are random variables correlated through an unmeasured normally distributed variable *U*, and depend linearly on a binary predictor *X*, inducing an effect on the means of *Y*_1_ and *Y*_2_. The conditional covariance between *Y*_1_ and *Y*_2_ can be expressed as *cov*(*Y*_1_, *Y*_2_|*X* = *x*) = *α*_1_*α*_2_*var*(*U*), and does not depend on *X*. Consider the alternative models: *Y*_1_ = *α*_1_*U* + *β*_1_*AX* + *ϵ*_1_ and *Y*_2_ = *α*_2_*U* + *ϵ*_2_, where *Y*_1_ and *Y*_2_ are correlated, and the variance of *Y*_1_ depends on the product of a latent continuous variable *A* multiplied by the binary predictor *X*. The conditional covariance can again be expressed as *cov*(*Y*_1_, *Y*_2_|*X* = *x*) = *α*_1_*α*_2_*var*(*U*), and does not depend on *X*. Finally, consider the models: *Y*_1_ = *α*_1_*UX* + *ϵ*_1_ and *Y*_2_ = *α*_2_*U* + *ϵ*_2_, where *Y*_1_ and *Y*_2_ are correlated, with the strength of the correlation depending on the predictor *X*. The covariance can now be expressed as *cov*(*Y*_1_, *Y*_2_|*X* = *x*) = *α*_1_*α*_2_*x var*(*U*), and does depend on *X*. Further details on those approximations are provided in the Supplementary Notes.

The approach can easily be extended to more than two outcomes by deriving an *N* × *p* matrix of products between centered outcomes, defined as **P** = *P*_1_, … *P*_*p*_, with *p*, the number of products, equals *k*(*k* − 1)/2 where *k* is the number of outcomes and *N* the sample size. The association between a predictor *X* and **P** can then be derived by applying a standard two-way analysis of variance (MANOVA), that is **P**∼**δ***X*. While valid, this approach is limited to situations where the effective sample size *N* is substantially larger than the number of products *p*. When this criterion is not met, we use principal component analysis to reduce the dimension of the product matrix, and use the top *m* principal components (PCs) to form an *N* × *m* matrix **Ω** used as input in our test. Given the independence between the principal components, we first considered using a sum of univariate PC tests to form a joint test, however, this approach was not calibrated (see ***Results*** and **Figs. S1-2**). Instead, we used a MANOVA, that is **Ω**∼**β***X*. For fast computation, the joint effect estimates 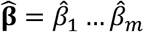 of association between each principal component *Ω*_*i*_ *i* ∈ [1 … *m*] and the predictor *X* are first derived using a single matrix operation: 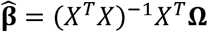. The Wilks’ lambda statistics, *W* = det(**Ω**^*T*^**Ω** − ***β****X*^*T*^*X****β***^*T*^)/det(**Ω**^*T*^**Ω**) is derived in a second step. Under the null hypothesis of no association, *W* follows a Fisher distribution *F*(*m, N* − *m* − 1). **Figure 1** presents an overview of the steps for applying the approach.

**Figure 1.**
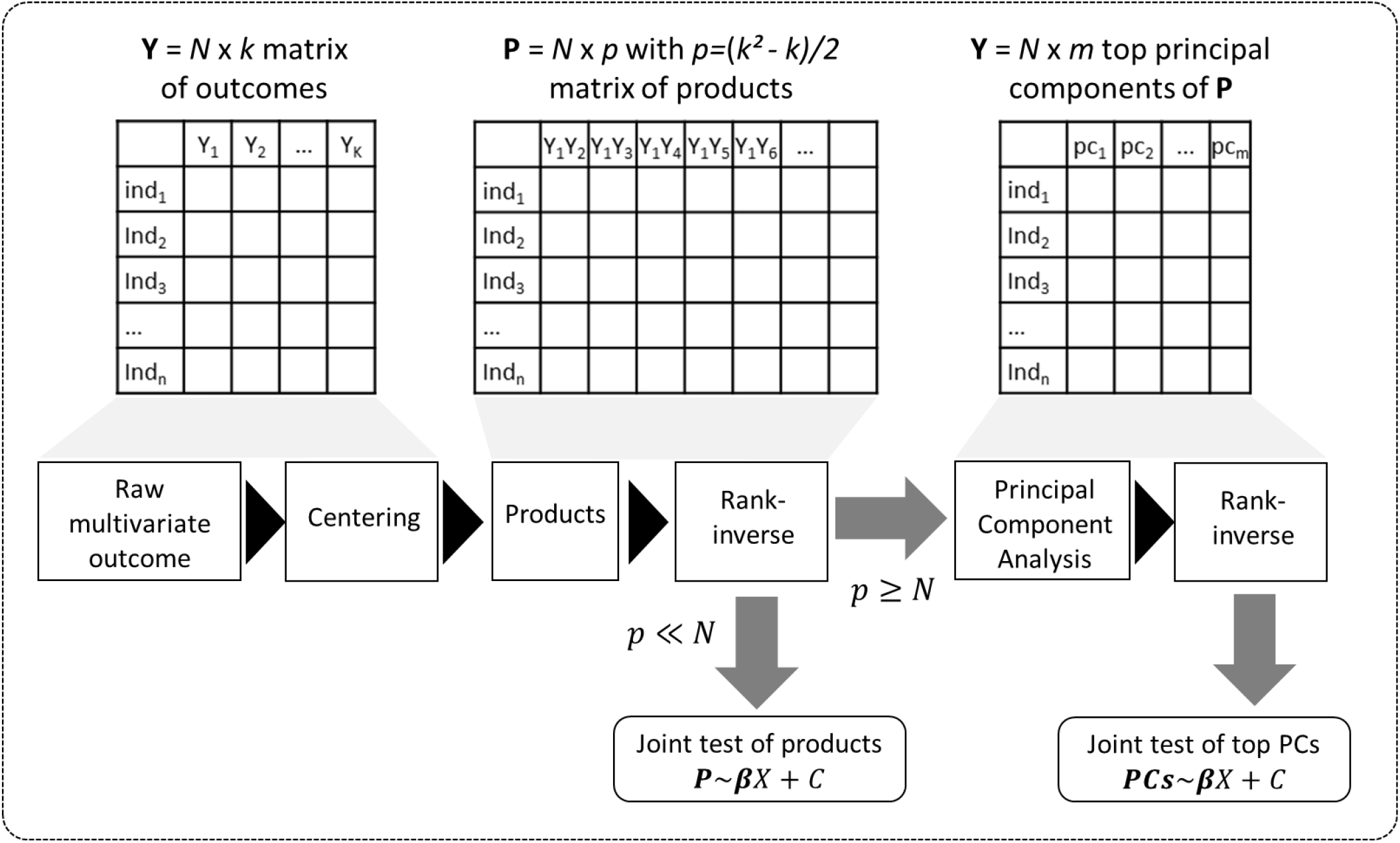
Overview of the MANOCCA approach. Starting with a multivariate outcome matrix of *N* samples and *k* variables, the data are first centered. The pairwise product of each of the *k* outcomes is computed, generating a high dimensional matrix of size *N* × *k*(*k* − 1)/2. If *N* >> *k*(*k* − 1)/2, a joint test of all products can be derived, otherwise the dimension of the product matrix is reduced using a Principal Component Analysis, to form a principal component space of size *N* × *m*. The final test, including covariates, can be performed on the products or the top *m* PCs using a Wilk’s lambda test.

In a standard MANOVA, potential confounding factors **C** = (*C*_1_ … *C*_*c*_) can be incorporated as covariate: **Ω**∼**C** + **β***X*. Again, for fast computation, we used a two steps procedure that consists in adjusting a priori both the outcome and the predictor for the covariates: 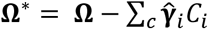, where 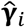 is a vector of estimated effect of *C*_*i*_ on **Ω**, and 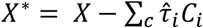, where 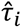 is the estimated effect of *C*_*i*_ on *X*, and applying the MANOVA on the residual variables: **Ω**^∗^∼**β***X*^∗^.

### Type I error rate simulation

We first assess the calibration under the null of MANOCCA and four existing approaches, Mantel test^11^, the Fisher method^12^, the Jennrich test^13^, and the BoxM test^14^, using fully simulated data and simple scenarios (**Fig. 2**). We drew a series of 10,000 replicates with a sample size of 1,000, each including a multivariate outcome **Y** and a binary predictor *X*∼*B*(0.4) drawn independently of **Y** under two different models. Note that we used a binary predictor as the four existing approaches do not allow for the analysis of continuous predictors. In the first model, replicates included five outcomes drawn from a multivariate normal with modest pairwise correlation. In the second model, replicates included 30 highly correlated non-normal outcomes drawn from a multivariate chi-squared distribution. The overall calibrations of all tests were derived by testing for association between *X* and the correlation between **Y** variables, and conducting a visual inspection of the *P*-value distribution.

**Figure 2.**
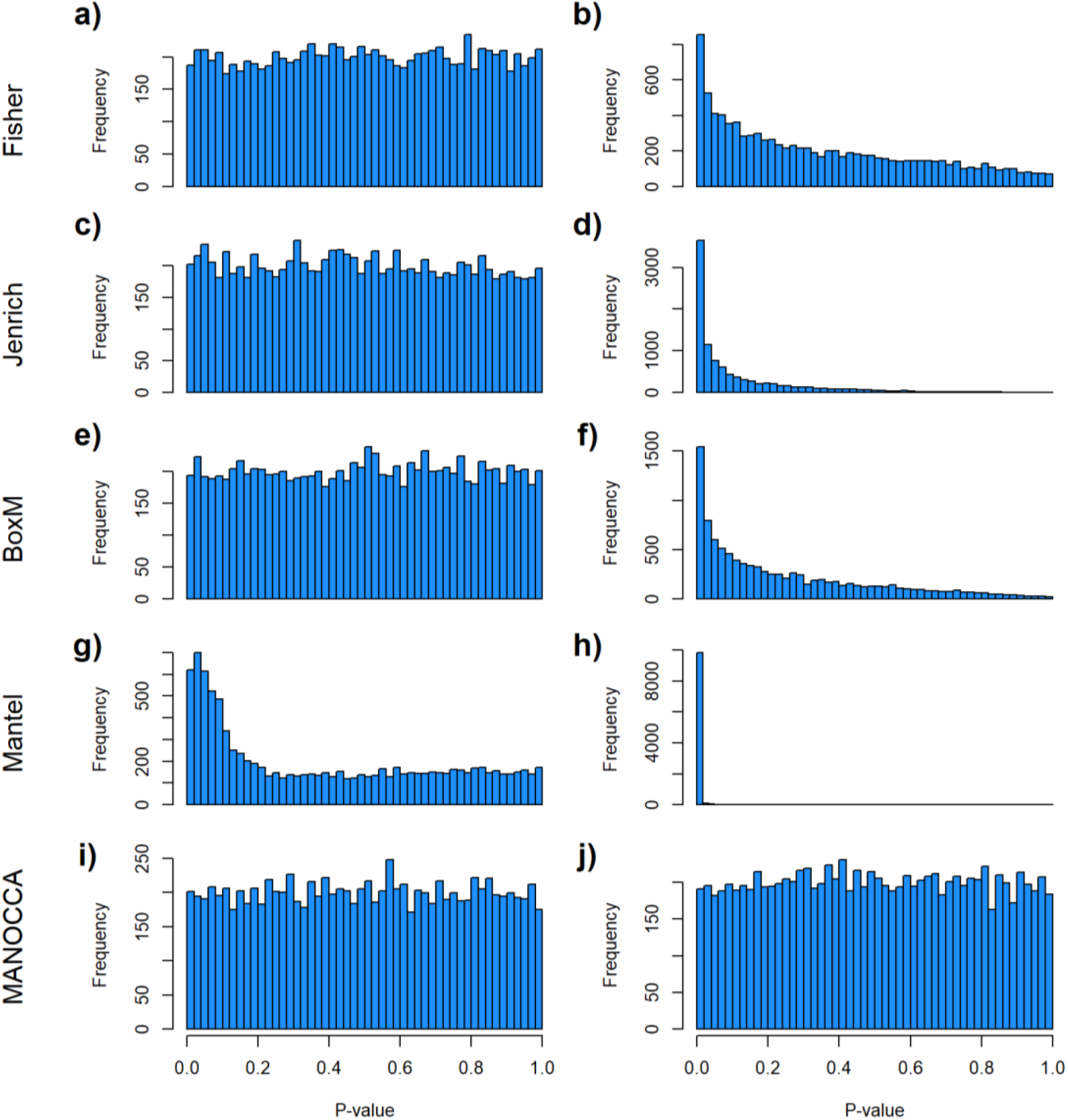
Limitation of existing methods. We assessed the calibration under the null of four approaches representing the state-of-the-art for covariance matrix comparison: the Fisher method (a,b), the Jenrich test (c,d), BoxM (e,f), and Mantel test (g,h), against the proposed MANOCCA approach (i, j). Note that we applied MANOCCA directly on the product matrix thanks to the high sample size compared to the number of products. We simulated series of 10 000 replicates, with a sample size of 1000 each, under two different null models. In the first model (a,c,e,g,i), replicates included five outcomes ***Y*** drawn from a multivariate normal with modest pairwise correlation. In the second model (b,d,f,h,j), closer to the expected distribution of omics data, replicates included 30 non-normal outcomes with high correlation. Calibration was derived by splitting each replicate in two random sets according to a random binary variable *X*∼*B*(0.4), and testing for association between *X* and the correlation between ***Y*** variables. The panels present the distribution of the *P*-values, expected to be uniformly distributed under this null model, for the five approaches and the two models.

We next assessed the robustness of MANOCCA under a wider range of scenarios, while modifying some of the modelling parameters. We first compared performance when using a binary or a continuous predictor (**Fig. S3**). We simulated a series of 100 replicates, each including 1,000 individuals and a multivariate outcome **Y** including 400 variables drawn from a multivariate chi-squared distribution with a point mass at 0 including 0% to 50% of the data. For each replicate we drew 1,000 predictors, either from a normal or binary distribution, and applied MANOCCA while varying the number of PCs used and applying no transformation, a rank inverse normal transformation on the product matrix, the principal component matrix, or both. The validity of MANOCCA was assessed using a Kolmogorov–Smirnov test for deviation from a uniform [0,1] distribution of the *P*-values across the 1,000 predictors tested. We then conducted simulations using real covariance matrices derived from the Milieu Interieur 169 flow cytometry-based variables and for a ranging sample sizes from 1000 until 5000 to draw guidelines on the parametrization of MANOCCA (**Fig. S6-7**). For predictors, we considered binary predictors with frequencies in [0.01 ; 0.40], but also categorical ones mimicking genetic variants with minor allele frequency in [0.01 ; 0.40], both generated independently of the multivariate outcome **Y**.

### Power simulation

To investigate power, we drew a series of 50 replicates with sample size of 1,000 including a binary exposure *X* with frequency of 0.5 and a multivariate outcome **Y** including 50 to 169 variables (**Fig. 4**). For each replicate, we used two covariance matrices, one for the exposed (*C*_1_), and the other for the unexposed (*C*_2_), and tested the association between *X* and **Y** using MANOCCA. We generated the outcome from a multivariate normal and real covariance matrix (*C*) derived from the Milieu Interieur flow cytometry data under three scenarios. In scenario (i), *C*_1_ = *C* and *C*_2_ = |*C*|^*γ*^ × *sign*(*C*), inducing a covariance with similar pattern among exposed but variability in the magnitude of covariance. In scenario (ii), *C*_1_ = *C* and *C*_2_ = *δC* + (1 − *δ*)Δ, where Δ is a random covariance generated using the R *randcorr* package^16^, thus inducing random noise between exposed and unexposed. In scenario (iii), we first drew *C*_1_ = *C*_2_ = *C* and then attenuated the covariance in an arbitrary chosen subset *ω* of *C*_2_, so that *C*_2{*ω*}_ = |*C*_{*ω*}_| ^*θ*^ × *sign*(*C*_{*ω*}_). We arbitrarily set *γ* to 1.5, *δ* to 0.2, and *θ* to 0.5, as it allowed for a similar average power across scenarios given the other simulation parameters.

### MANOCCA association screening in Milieu Intérieur

The *Milieu Intérieur* (MI) Consortium is a population-based cohort initiated in September, 2012^17^. It comprises 1,000 healthy volunteers from western France, with a 1:1 sex ratio. The cohort collected a broad range of variables, including genomic, immunological, environmental, and clinical outcomes. We conducted systematic MANOCCA screenings for environmental effects on the covariance of two sets of data: 169 flow cytometry-based immune cell phenotypes^18^ and 33 health-related blood biomarkers, including 22 metabolites and 11 cell counts^17^ (**Table S1**-**S2**). We focused on two types of predictors: health and lifestyle factors collected from questionnaires, and genome-wide variants. Health and lifestyle factors included demographics, medical and vaccination history, psychological traits, socio-professional information, smoking habits, physiological measurements and nutrition measured as part of the Nutrinet^19^ study (**Table S3**). After the filtering of ancestral outliers individuals^20,21^, the genetic screening was conducted in 894 participants for a total of 5,667,803 variants after filtering and imputation using IMPUTE2^22^. Except when used as predictor, all analyses were adjusted for age, sex and body mass index (BMI). For blood metabolites, the number of products allowed for a direct analysis of the products without requiring the PCA step, and we considered both the products and the PCs as outcomes. For comparison purposes, we also conducted, for each screening, a standard MANOVA on the mean of the multivariate outcome (see **Supplementary Notes**).

#### Human samples

Samples came from the Milieur Intérieur Cohort, which was approved by the Comité de Protection des Personnes – Ouest 6 (Committee for the protection of persons) on June 13th, 2012 and by French Agence nationale de sécurité du médicament (ANSM) on June 22nd, 2012. The study is sponsored by Institut Pasteur (Pasteur ID-RCB Number: 2012-A00238-35), and was conducted as a single centre interventional study without an investigational product. The original protocol was registered under ClinicalTrials.gov (study# NCT01699893). The samples and data used in this study were formally established as the Milieu Interieur biocollection (NCT03905993), with approvals by the Comité de Protection des Personnes – Sud Méditerranée and the Commission nationale de l’informatique et des libertés (CNIL) on April 11, 2018. All donors gave written informed consent. All data used in this study are available at https://dataset.owey.io/.

## Results

### Method comparison and MANOCCA characteristics

We identified four existing approaches allowing to test for the effect of a predictor on the covariance matrix of a multivariate outcome: (i) the Mantel test^11^, which consists in deriving a distance metric between two square matrices of the same dimension and comparing this distance to an empirical distribution derived through permutation; (ii) the Fisher method^12^, which builds a statistic based on the sum of the squared correlations over all cells from the covariance matrix; (iii) the Jennrich test^13^, which, in its simplest form, consists in estimating the statistic based on the Hadamard product of a given correlation matrix and the inverse of a second matrix of the same dimension; and (iv) the BoxM test^14^, which extends the Levene’s test of homogeneity of variance, an approach often used in human genetics^23^. Further description of each of the four approaches is provided in **Supplementary Notes**. We conducted series of simulations to assess their robustness under the null using a binary predictor and no covariates, as these approaches cannot handle continuous predictors and do not allow adjustment for covariates. Except for the Mantel test, all methods performed relatively well for a simple model with a few normally distributed outcomes. Conversely, they all displayed severe type I error rate inflation when confronted with non-normal correlated variables, mimicking omics data (**Fig. 2a-h**). In comparison, when applied to the same simulated data, MANOCCA was correctly calibrated in all simulations (**Figure 2i-j**).

The effect of the predictor on the covariance, the mean and the variance of a set of outcomes are expected to be statistically independent (see **Supplementary Notes**). We confirmed this orthogonality between mean (derived using a two-ways MANOVA), variance (Levene’s test), and the proposed covariance tests through simulation. **Figure 3** shows that, under realistic modelling assumptions, the MANOCCA test captures only effects on the covariance, and can therefore identify effects missed by both mean and variance-based approaches. **Figure 3d** further illustrates bivariate data where a binary predictor *X* is associated with covariance but neither the mean nor the variance of the outcomes. Importantly, the independence of the three tests does not imply signal across the three approaches will necessarily be uncorrelated in real data. Indeed, one can easily draw scenarios with *e*.*g*. effect of a predictor on both the mean and covariance of a multivariate outcome. Also, unless specified otherwise, we modelled the effect of a predictor on the covariance through an interaction with a latent variable associated shared across the outcomes tested (see ***Methods***). Under this modelling, effects on the covariance can in general be transposed to effects on the correlation. However, when the predictor has an effect on the variance of either outcome, this equality is not valid anymore, as the correlation will depend on *X*, while the covariance will not (**Supplementary Notes**).

**Figure 3:**
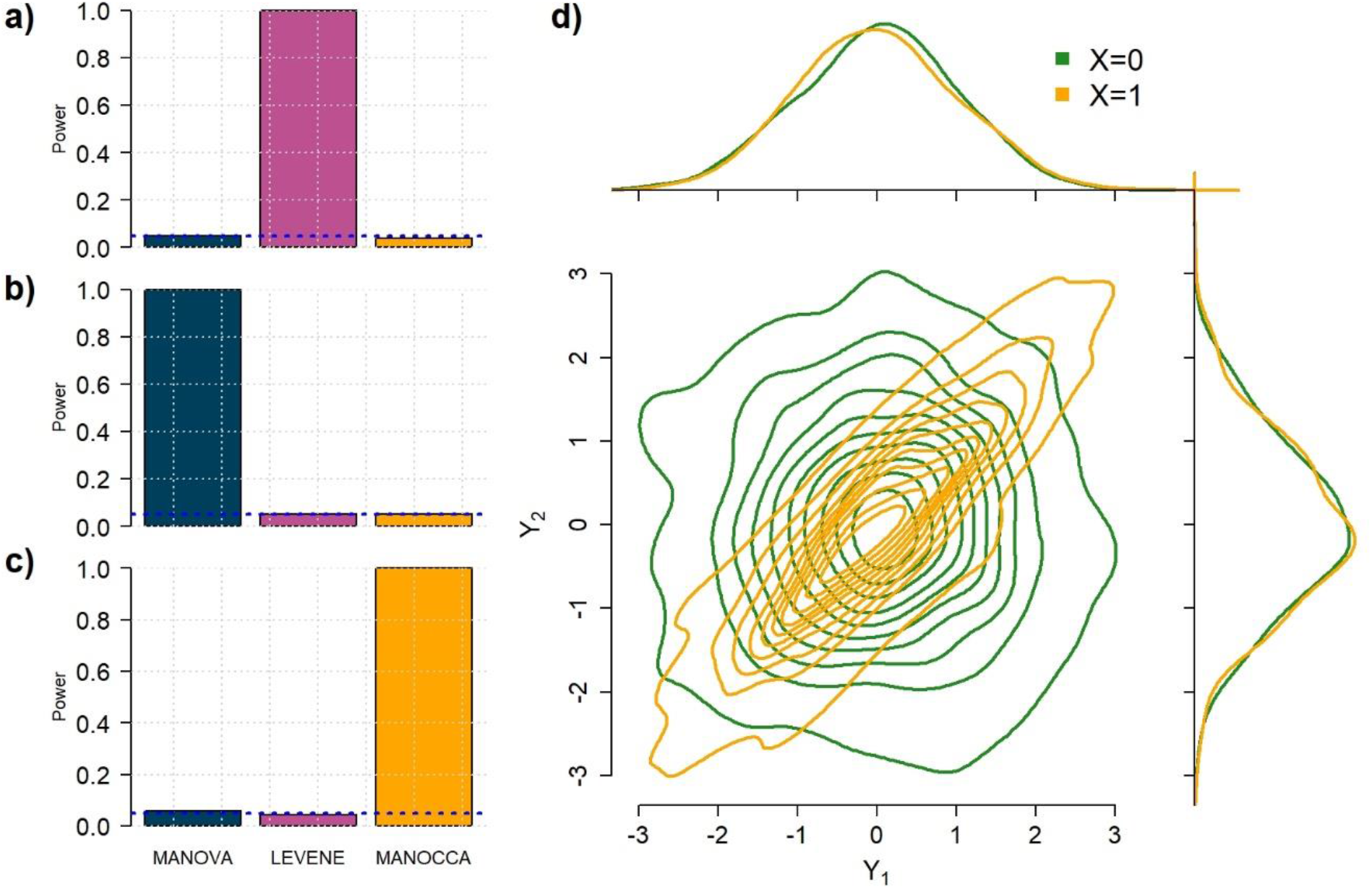
Orthogonality between MANOCCA and other tests. We simulated series of datasets under three models where a binary predictor *X* influences orthogonally either the mean, the variance or the covariance of a bivariate outcome ***Y***. In model a), each outcome *Y*_*i*_ is drawn from a standard additive model: *Y*_*i*_ = *U* + *β*_*i*_*X* + *ε*_*i*_, where *U* is a normally distributed variable shared across *Y*. and *ε*_*i*_ are independent normal residuals. In model b), each *Y*_*i*_ is drawn from *Y*_*i*_ = *U* + *γ*_*i*_*A*_*i*_*X* + *ε*_*i*_, where *A*_*i*_ is normally distributed variables producing heterogeneity in the variance of *Y*_*i*_ conditional on *X*. In model c), each *Y*_*i*_ are drawn from the interaction model *Y*_*i*_ = *δ*_*i*_*UX* + *ε*_*i*_, that produces heterogeneity in the correlation across *Y*_*i*_ conditional on *X*. For each model, we derived the power at the *P*-value threshold of 0.05 for a joint mean effect test (MANOVA), a test of variance for a randomly selected *Y*_*i*_ (LEVENE), and the proposed covariance test (MANOCCA). The parameters *β*_*i*_, *γ*_*i*_ and *δ*_*i*_ were chosen to maximize the power of the at least one of the three tests. The blue dash line indicates the *P*-value threshold of 0.05. Panel d) shows an example of a bivariate distribution where *X* is not associated with the mean and variance of the two outcomes but with their covariance.

**Figure 4.**
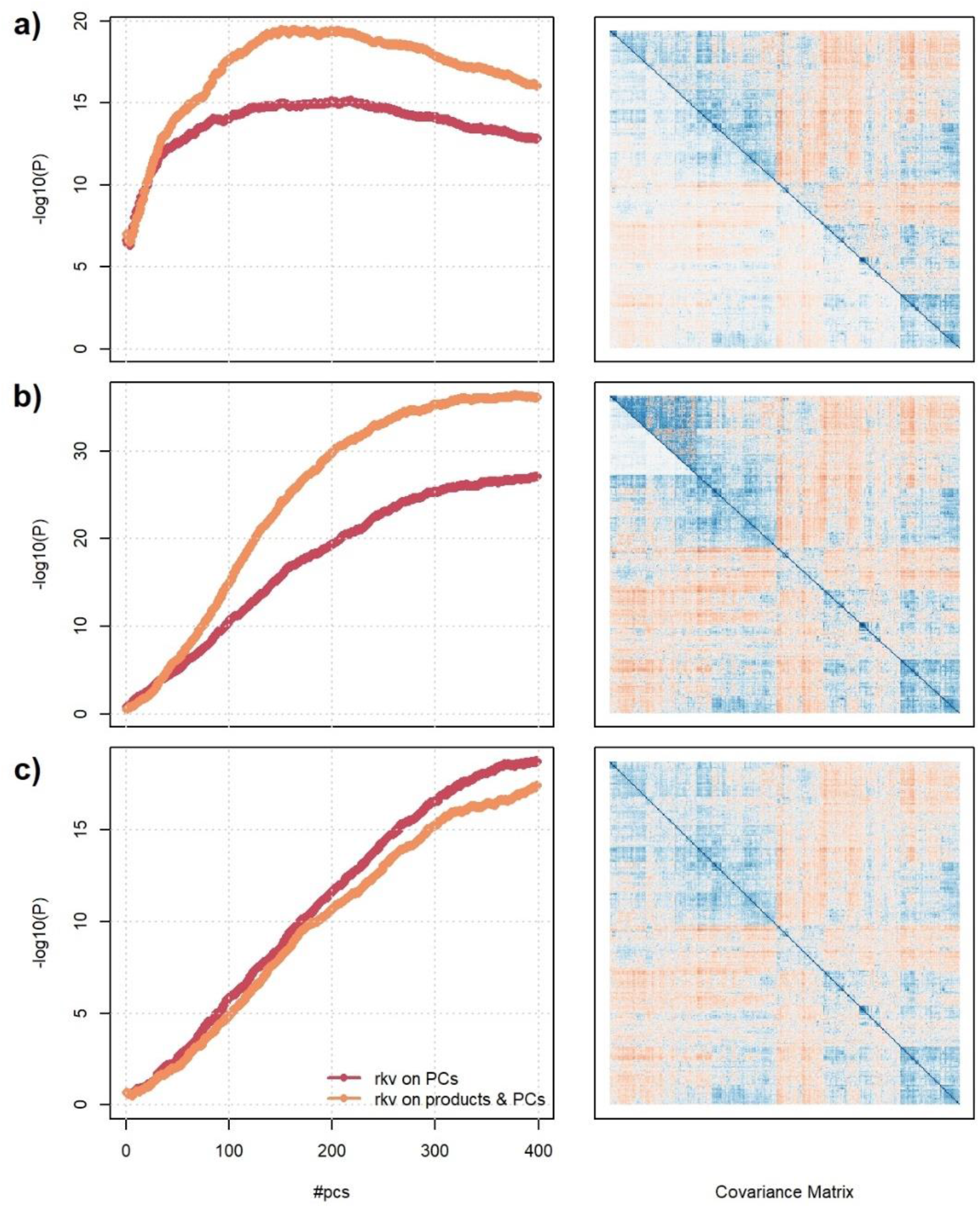
Impact of transformation and number of PCs on statistical power. Power of MANOCCA as a function of the number of principal components retained in the joint test (2 to 400), while applying two different pre-processings: a rank-inverse normal (rkv) on the principal component only, or a rank-inverse normal on the principal components and the products. We drew series of 50 replicates with sample size 1,000, including a binary exposure with frequency of 0.5 and 30 to 169 outcomes. For each replicate, we drew two covariance matrices, one for the exposed, and one for the unexposed. We generated the outcome under three scenarios using a multivariate normal and covariance derived from real data. In panel a) the two matrices are similar but with attenuated covariances among exposed. In panel b) random noise is added to the covariance of the exposed group. In panel c) the two matrices are equal, except for a subset of outcomes where the covariances have been attenuated. Left panels show the average over the 100 replicates of the -log_10_(*P*-value) derived using MANOCCA for the two pre-processings. Right panels present the matrix produced for each scenario using data from an arbitrarily chosen replicate, with upper and lower triangles showing the true covariance in unexposed and exposed, respectively.

### Extension to high-dimension data

MANOCCA is readily applicable to the matrix of outcome product in all scenarios where *p* is substantially smaller than *N*. We used Principal Components Analysis (PCA) to reduce the dimension of *P*, using the top principal components as the primary outcome. As for all linear models, the maximum number of PCs that can be analysed jointly remained bounded by the sample size. More generally, high dimension outcomes data brings the question of the latent space dimension, that is, the number of PCs kept in the analysis to achieve maximum power while maintaining a correct type I error rate. Moreover, by construction, both products and PCs tend to display kurtotic distributions, especially for omics-like data, which might also impact performance. We conducted a series of simulations to investigate the validity of these two components. Specifically, we measured the type I error rate while varying the number of top *m* PCs selected, and applying (i) no transformation, or an inverse rank transformation on (ii) the product, (iii) the PCs, or (iv) both products and PCs. As shown in **Figure S3a**, if the predictor being tested is continuous, the test remains well calibrated regardless of the transformation applied, allowing for the use of a large number of PCs. Conversely, when analysing binary predictors, the test requires a normalization of the PCs, only allowing a limited number of PCs to be analysed jointly (**Fig. S3b**). **Figure S4-5** present the results from extended simulations, providing guidelines to determine the number of PCs that can be analysed jointly conditionally on the predictor frequency and the cohort sample size.

We next evaluated the power of MANOCCA across different scenarios in which the true covariance depends on a binary predictor *X* with a frequency of 0.5. We tested up to 400 PCs, and normalized either the products and PCs or the PCs only, the two transformations that display a calibrated null distribution (**Fig S3**). The optimal number of PCs varied substantially across the simulated scenarios. **Figure 4** illustrates three complementary cases. When *X* acts as a global scaling factor of the covariance, the maximum power is observed when using a limited number of top PCs and decreases after reaching that optimum (**Fig. 4a**). When *X* affects only a subset of outcomes, the maximum power is reached when including a fairly large number of PCs and converge afterward (**Fig. 4b**). As expected, when *X* induces random noise in the covariance matrix, the power increases continuously with the number of PCs (**Fig. 4c**). Among the scenarios we considered, the double normalization (on products and PCs) produced on average larger power, and this transformation was therefore used in all subsequent analyses.

### Efficient implementation

The MANOCCA approach requires multiple steps that can be computationally expensive in large-scale data. The main limiting step is the computation of the product matrix followed by the PCA transformation, but it is a one-time cost regardless of the number of predictors tested. With *N* the sample size, *k* the number of outcomes and *q* the number of predictors to test, the computation time is divided in *O*(*Nk*^2^) for the computation of the product matrix, *O*(*max*(*N, k*^2^)^2^ ∗ *min*(*N, k*^2^)) for the computation of the PCA, and *O*(*Nq*) for the test of *q* predictors (**Fig. 5**). Most steps were implemented with limited usage to exterior libraries, but ground proofed against multiple existing tools. The approach is implemented in a *Python* package with dependencies to *numpy, scipy* for the fisher distribution, *scikit-learn* for the PCA and *pandas* for *dataframe* integration, but all computations are performed under *numpy* array to increase performance. Each step was optimized to minimize computational time and, given that most steps are independent, especially the product matrix, the python version allows for a user-friendly parallel computing implementation if multiple cores are available. An R version, though less optimized, is also available and was used to verify our results.

**Figure 5.**
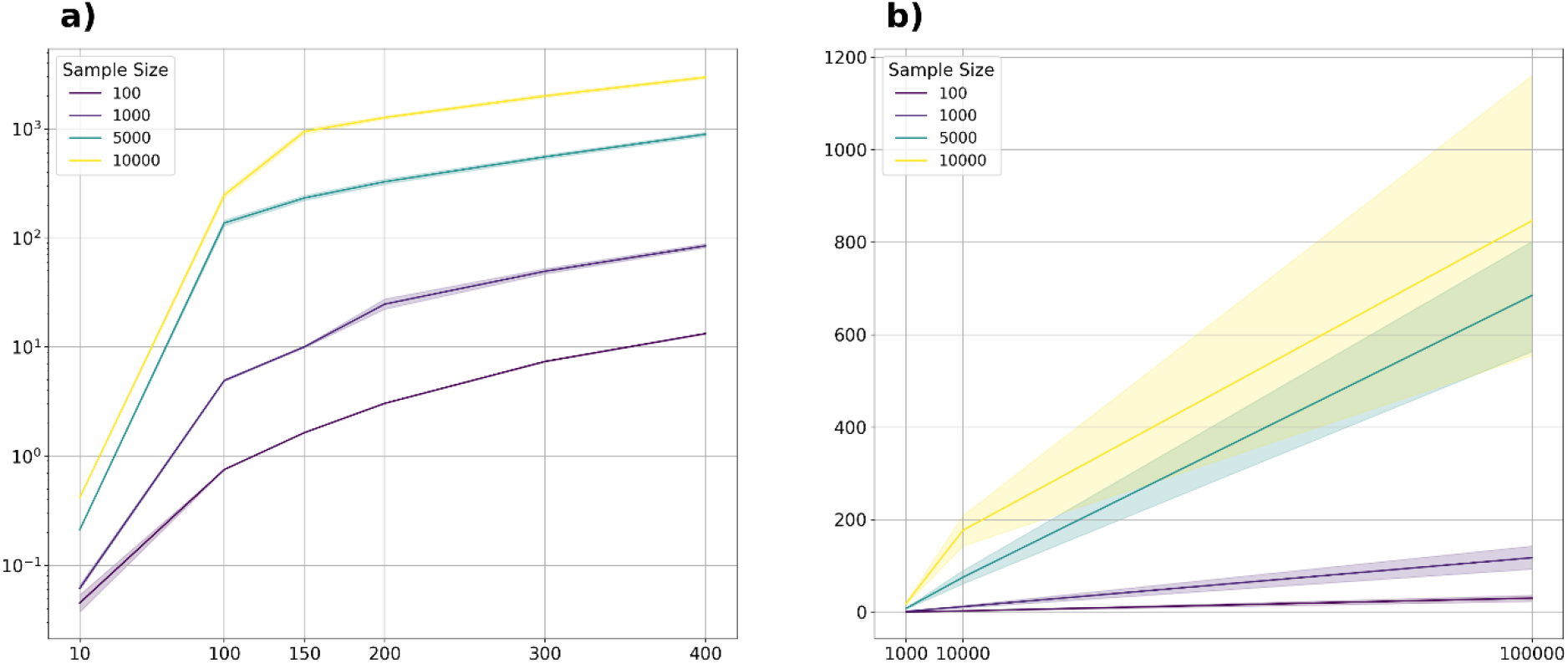
Computational efficiency. Computational time for running MANOCCA on varying sample sizes, numbers of outcomes and numbers of predictors tested. Over 1000 simulations each time, we simulated a multivariate normal distributions of various sizes and computed the product Matrix > Rank transform > PCA > Rank transform using sequential computation. Panel a) displays the running time to transform the outcome matrix into the reduced covariance matrix for the test, as a function of the number of outcomes (10, 100, 150, 200, 300, 400), and sample size (100, 1000, 5000 and 10000). Y-axis is in log_10_-scale. Panel b) displays the running time for the testing part for a ranging number of sample sizes (100, 1000, 5000 and 10000) and ranging number of predictors (1000, 10000, 100000).

### Application to Milieu Intérieur omics data

We applied MANOCCA in 1,000 healthy individuals from the *Milieu Intérieur* (MI) cohort to screen for factors associated with changes in the covariance of 33 blood biomarkers and 169 flow cytometry-based immune phenotypes (**Table S1-3**). Both datasets display high correlation, ranging from -0.71 to

0.99 for the flow cytometry data, and from 0.08 to 0.98 for the biomarkers (**Fig. S6**). We first investigated the effect of 49 health-related and lifestyle factors using a subset of 992 participants with complete data. We applied the proposed PCA reduction, and investigated power when using 5 to 200 top PCs with a step of 5, resulting in 40 tests per variable. As showed in **Figure 6a-b**, multiple features were associated at a stringent Bonferroni corrected significance level (*P* < 2.5 x 10^-5^ accounting for 1,960 tests). Flow cytometry-based phenotypic covariance was associated with age (*P* < 6.0 x 10^-9^) and all smoking variables: smoking status (*P* < 2.1 x 10^-12^), smoking frequency (*P* < 1.2 x 10^-11^), number of years smoked (*P* < 3.8 x 10^-10^), number of years since last smoke (*P* < 7.5 x 10^-9^). Likewise, blood biomarkers covariance was strongly associated with age (*P* < 3.5 x 10^-33^) and sex (*P* < 1.3 x 10^-30^), and to a lesser extent with BMI (*P* < 1.6 x 10^-7^) and smoking variables (minimum *P* < 5.5 x 10^-12^, for smoking status). Except for the age-flow cytometry association, the maximum association signal was almost reached when including the top 50 PCs and display only modest improvement when including more PCs (**Fig. 6**).

**Figure 6.**
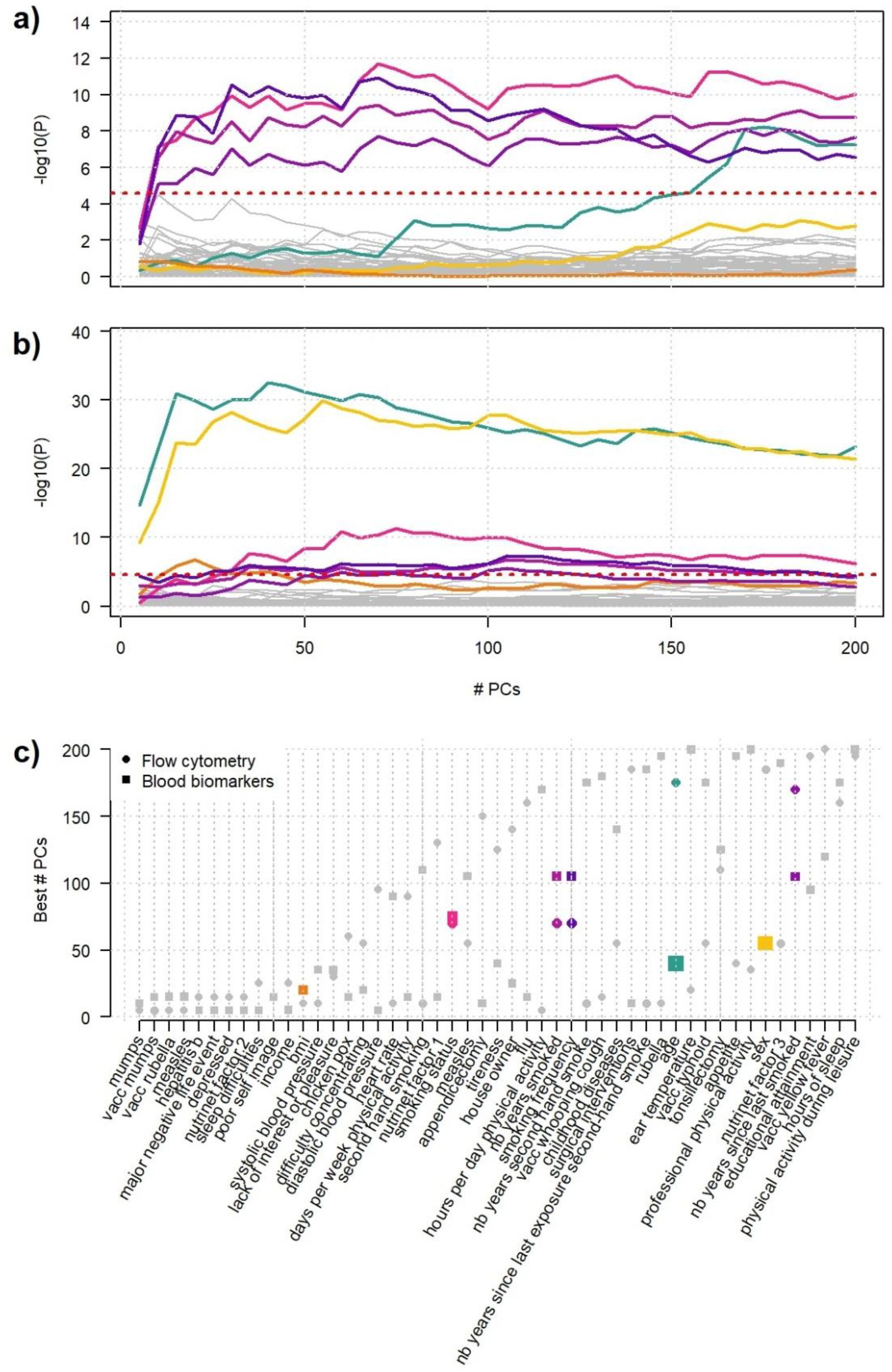
Screenings for host factors. Screenings for effect of 49 health and lifestyle factors on the covariance of 169 flow cytometry-based immune phenotypes (a) and 33 blood biomarkers (b) using the MANOCCA approach. We ran each screening using 5 to 200 principal components (PC) with a step of 5. Variables with a *P*-value below the adjusted Bonferroni threshold (2.6 x 10^-5^, red dashed line) for each screening are displayed in colour: age (green), sex (yellow), body mass index (orange), smoking (purple gradient). Panel c) presents the list of predictors considered and displays the number of PCs corresponding to the minimum *P*-value observed.

Thanks to a limited number of outcomes, the blood biomarker dataset could also be analysed by applying MANOCCA directly on the 528 pair-wise products. To investigate the value of using the PCA in such situations, we applied MANOCCA on the product and compared results against the PCA-based approach. Note that the product-based test should be approximately equivalent to the test of all PCs, which was confirmed for these data (**Fig. S7**). Comparing the minimum *P*-value across the 40 PCA-based test (*P*_*PCA*_) against the *P*-value from product-based test (*P*_*prod*_), we observed a substantial gain for the PCA-based approach, even when accounting for the multiple testing cost of the PCA approach. The product-based test identified only five of the seven signals from the PCA-based approach (**Fig. 7**). For the strongest signals, the association *P*-value from the best PCA-based test was several orders of magnitude larger than for the product-based test (*e*.*g*., for age, *P*= 3.5 x 10^-33^ and 2.6 x 10^-11^, respectively). This suggests that the benefit of testing multiple sets of PCs can strongly outpace the statistical cost of multiple testing.

**Figure 7.**
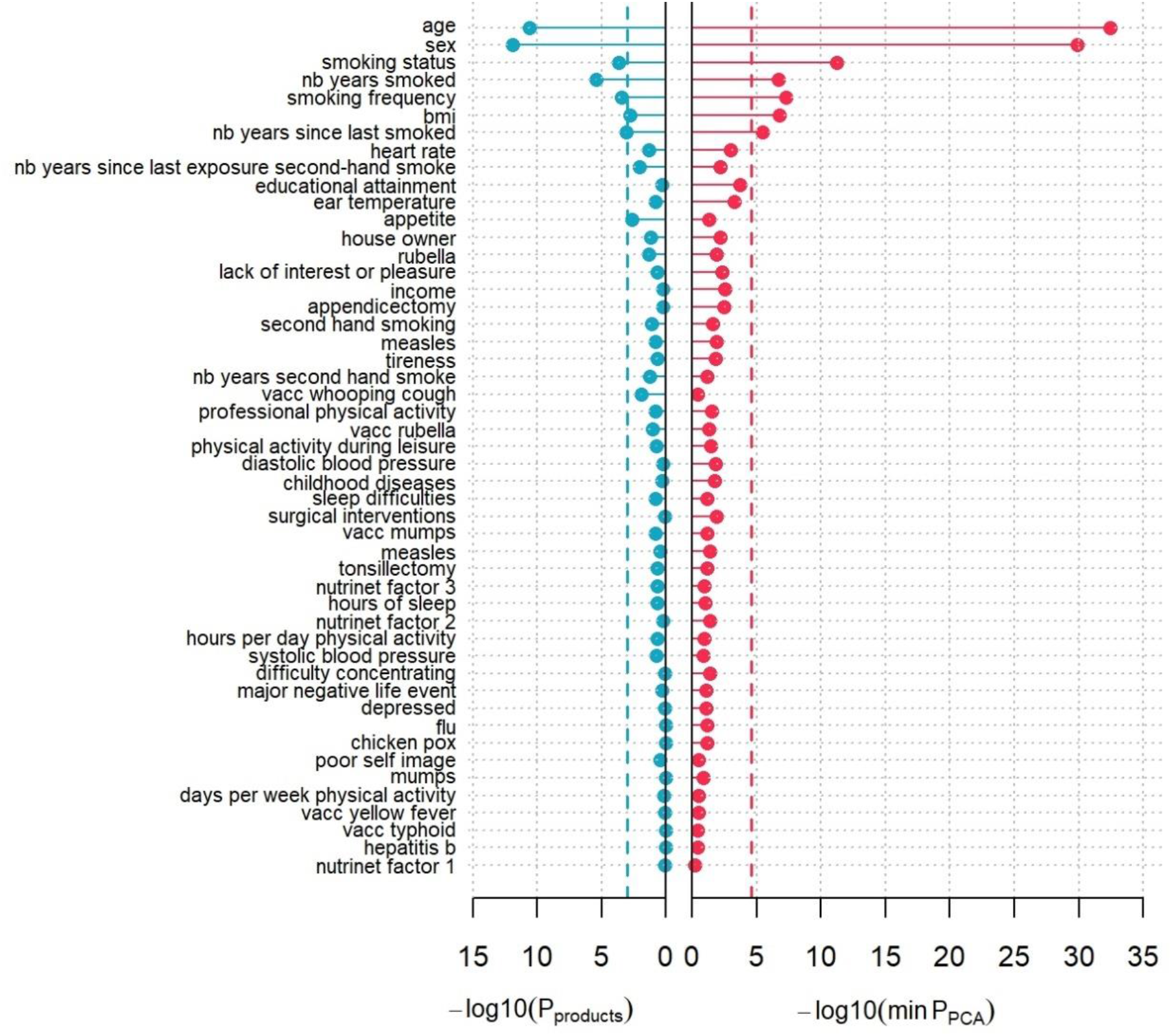
Comparison of PCA-based and product-based MANOCCA. Screening comparison for covariance signal using MANOCCA on the 33 blood biomarkers as outcomes and using the health and lifestyle eCRF questionnaire data as predictors. We ran the MANOCCA screening using the full 528 pair-wise biomarkers products (blue dots) and the principal components (PCs) derived from the product matrix (red dots). For the latter, we kept the min(*P*-value) out of 40 models including 5 to 200 PCs with a step of 5 PCs. The corresponding Bonferroni correction threshold (dashed line) were derived for each approach based on the number of tests conducted (*P-value* threshold of 1.0×10^-3^ and 2.6×10^-5^, respectively).

When comparing the MANOCCA results to a standard MANOVA applied to both datasets, MANOVA identified significant associations (*P* < 1.0 x 10^-3^ to account for the 49 tests conducted) with 8 and 13 predictors associated in the flow cytometry data and the blood biomarkers data, respectively (**Table S4**). These associations included all associations detected by MANOCCA. More generally, while the two tests are expected to be independent, we observed a strong correlation between association signal as measured by the -log10(*P*-value) (*ρ*_*biomarkers*_ = 0.88, *ρ*_*cytometry*_ = 0.65), suggesting that many of the predictors are associated with effects on both the mean and variance of the outcome studied (**Fig. S8**). However, several predictors display discordant associations, with a significant effect on the mean but no effect on the covariance. This includes, for example, systolic blood pressure (*P*_*Manova*_ = 2.6 x 10^-11^, *P*_*MANOCCA*_ = 0.14), and heart rate (*P*_*Manova*_ = 4.9 x 10^-24^, *P*_*MANOCCA*_ = 0.001) effect on blood biomarkers. On the other hand, one predictor, the number of years of second-hand smoking, display suggestive significance with MANOCCA on the flow cytometry data, but did not display mean effect (*P*_*Manova*_ = 0.31, *P*_*MANOCCA*_ = 3.0 x 10^-5^).

Finally, we conducted genome-wide association studies (GWAS) for both blood biomarkers and flow cytometry datasets, testing 5,699,237 genetic variants with a minor allele frequency (MAF) > 5% in up to 894 samples where both genetic and phenotypic data were available. All tests were adjusted for age, sex, BMI and the 10 first genetic PCs of the genotyping matrix. Note that for this genetic screening we only applied the adjustment on the outcomes. Two-sided adjustment has already been used for mean effect tests in GWAS to account for relatedness^20^, but would require further investigation to be extended in the MANOCCA test. Following the results from our simulations for genetic variants with a MAF 5% or larger, the type I error will remain robust only for up to 50 principal components (**Fig. S4c**), resulting in 10 GWAS per dataset. MANOCCA did not detected any genome-wide significant signals at a stringent Bonferroni corrected threshold (*P* = 5 x 10^-9^, **Fig. S9**). Yet, 46 genetic variants from eleven loci show suggestive significance association (*P* = 5 x 10^-7^, **Table S5**). We conducted phenome-wide association study on each variants using the *ieu* database API^24^. Most variants showed strong association with multiple phenotypes from this database (**Table S6**). In particular, four out of the eleven loci harboured genetic variants that were eQTL for one or multiple genes (**Table S7**), suggesting that our covariance-based approach might capture variants involved in the regulation of gene expression.

## Discussion

Covariance is a fundamental feature of omics data. Covariances might be explained by multiple factors, including shared biological mechanisms, shared environmental risk factors, or causal effects between the outcomes measured. However, our understanding of the factors involved in covariance has been very limited, partly due to the lack of adapted methodologies and software allowing for systematic screening of large-scale omics datasets. Here, we present MANOCCA, a robust and computationally efficient approach for the identification of predictors associated with the covariance of a multivariate outcome. We show that MANOCCA outperforms existing covariance methods and that, given the appropriate parametrization, it can maintain a calibrated type I error in a range of realistic scenarios when analysing highly multidimensional data. The application of MANOCCA to the *Milieu Intérieur* dataset demonstrates the validity and relevance of our approach, identifying multiple health-related and lifestyle factors significantly associated with the covariance of blood biomarkers and immune phenotypes.

The MANOCCA approach has three main limitations. First, we used principal components analysis to address situations where the number of covariance terms is larger than the sample size. We defined guidelines that constrain the maximum number of PCs that can be used to ensure the validity of the test when analysing binary or categorical predictors. Regarding power, both simulation and empirical data analyses show that the optimal number of PCs to be included to maximize power can vary substantially conditional on the true covariance association pattern. Here, we use systematic screenings testing a range of PCs, and corrected the association results for multiple testing. Our analyses suggest that the benefit of this strategy largely overcomes its statistical cost. Note that this correction strategy might be further improved as it does not account for correlation between each PC test. Second, our extensive simulation analyses show that when reaching high dimensions, the validity of the test relies on a strong data pre-processing to circumvent the non-normal distribution of products and principal components. Future work is required to identify a refined modelling of these non-normal distributions and avoid this pre-processing. Third, we investigated the performance of our approach on two types of omics data (blood flow cytometry and metabolites), and confirmed its validity and power in these data. Omics data from other sources (e.g., RNAseq) might carry additional complexity that would have to be investigated by simulations before conducting real data applications.

Given the increasing number of high dimensional omics data available in existing human cohorts, our approach provides opportunities to investigate multivariate outcomes from a new perspective. Because MANOCCA is built on a standard linear framework, the approach can be extended in many directions, including the derivation of the individual contribution of outcomes and the development of predictive models. Altogether, we expect the application of our method to produce novel insights on the complex structure linking highly intertwined omics data.

## Supporting information

SuppTables

## Acknowledgments

This research was supported by the Agence Nationale pour la Recherche (ANR-20-CE15-0012-01). This work has been conducted as part of the INCEPTION program (Investissement d’Avenir grant ANR-16-CONV-0005).

The Milieu Intérieur Consortium¶ is composed of the following team leaders: Laurent Abel (Hôpital Necker), Andres Alcover, Hugues Aschard, Philippe Bousso, Nollaig Bourke (Trinity College Dublin), Petter Brodin (Karolinska Institutet), Pierre Bruhns, Nadine Cerf-Bensussan (INSERM UMR 1163 – Institut Imagine), Ana Cumano, Christophe D’Enfert, Ludovic Deriano, Marie-Agnès Dillies, James Di Santo, Gérard Eberl, Jost Enninga, Jacques Fellay (EPFL, Lausanne), Ivo Gomperts-Boneca, Milena Hasan, Gunilla Karlsson Hedestam (Karolinska Institutet), Serge Hercberg (Université Paris 13), Molly A Ingersoll (Institut Cochin and Institut Pasteur), Olivier Lantz (Institut Curie), Rose Anne Kenny (Trinity College Dublin), Mickaël Ménager (INSERM UMR 1163 – Institut Imagine), Frédérique Michel, Hugo Mouquet, Cliona O’Farrelly (Trinity College Dublin), Etienne Patin, Sandra Pellegrini, Antonio Rausell (INSERM UMR 1163 – Institut Imagine), Frédéric Rieux-Laucat (INSERM UMR 1163 – Institut Imagine), Lars Rogge, Magnus Fontes (Institut Roche), Anavaj Sakuntabhai, Olivier Schwartz, Benno Schwikowski, Spencer Shorte, Frédéric Tangy, Antoine Toubert (Hôpital Saint-Louis), Mathilde Touvier (Université Paris 13), Marie-Noëlle Ungeheuer, Christophe Zimmer, Matthew L. Albert (In Sitro), Darragh Duffy§, Lluis Quintana-Murci§,

¶ unless otherwise indicated, partners are located at Institut Pasteur, Paris

§ co-coordinators of the Milieu Intérieur Consortium

## Code availability

All code is available in Python and R at: https://gitlab.pasteur.fr/statistical-genetics/manocca

Additional information can be found at: http://www.milieuinterieur.fr

