## Supplementary materials for "A multivariate outcome test of covariance"

### Supplementary Material

### Supplementary Notes

#### *Overview of the state-of-the-art*

We identified four approaches allowing for the comparison of covariance matrices: the Fisher method[1], the Jenrich test[2], the BoxM test[3], and the Mantel test[4].

The Mantel test is a method that uses permutations to compare the similarity of two distance matrices. It is commonly applied in microbiology using Bray–Curtis dissimilarity matrices[5]. Consider two square distance matrices $D^x$ and $D^y$ of the same dimension $n \times n$, the Mantel test is derived using a three steps process:

1) Compute the matrix distance $r = \sum_{i=1}^{n} \sum_{j>i}^{n} D_{i,j}^{x} * D_{i,j}^{y}$
2) For a number of times permute the rows and columns and compute again the matrix distance $\tilde{r} = \sum_{i=1}^{n} \sum_{j>i}^{n} D_{i,j}^{x} * \widetilde{D}_{i,j}^{y}$
3) Derive the empirical *P*-value as the proportion of permutation where $r$ is larger than $\tilde{r}$

The Mantel test is fairly easy to implement and doesn't need any prior distribution hypothesis, it only works for positive distance matrices. It is also relatively slow compared to other methods as it is permutation based, and is restricted to a binary predictor. In our analysis, to match the data used in the other tests we applied it on the square of the covariance matrices.

Box M's test is a statistical method that generalizes the Bartlett test [6] for multidimensional settings. It is used to test the equality of multiple variance-covariance matrices. Consider a matrix $Y = (Y^1, \ldots, Y^P)$ following a multivariate normal distribution. Given $K \geq 2$ subsets of $Y$, derive the corresponding covariance matrices: $C_{Y_{k=1}}, \ldots, C_{Y_{k=K}}$. The Box M's test of homogeneity of variance and covariance between the $K$ matrices is defined as:

$$M = (n_K)\,\mathrm{logdet}(W) - \sum_{k=1}^{K} (n_k - 1)\mathrm{logdet}(C_{X_{y=k}})$$

where, $W = \sum_K \left[(n-K)C_{X_{y=k}}\right] / (n-K)$. Under the null the $M$ statistics follows a chi-squared distribution:

$$M(1-\delta) \sim \chi_{df}^2$$

where $\delta = \frac{2p^3+3p+1}{6(p+1)(K+1)} [(\sum_{k=1}^{K} \frac{1}{n_k-1}) - \frac{1}{n-K}]$, and $df = \frac{p(p+1)(K-1)}{2}$

The Box's M test is a non-parametric method which does not require prior knowledge of the underlying data distribution and is rather robust to violations of assumptions such as normality and homogeneity of variance and covariances. On the downside it can be rather computationally heavy to compute and do not account for multiple comparisons meaning the type I error rate will increase when comparing multiple matrices. It is also known to be biased when the sample size is too small.

Fisher Z-score test is a statistical method proposed by Steiger[1] that is used to test the null hypothesis that the sum of squared correlations or the sum of squared Fisher transformed correlations is distributed as a chi-squared distribution. The test statistic is calculated by summing the squared Fisher Z-transformed correlation coefficients and comparing the resulting value to a chi-squared distribution to determine the significance of the correlation. With $C_1$ and $C_2$ two correlation matrices derived from datasets of sample size $n_1$ and $n_2$, the test can be defined as followed :

$$F = \left(\frac{n_1 n_2}{n_1 + n_2} - 3\right) \sum_i \sum_{i<j} \left(C1_{ij} - C2_{ij}\right)^2 \sim \chi^2_{n_1 n_2}$$

The correlation coefficient of the correlation matrices can also be transformed using the Fisher transformation: $f(x) = \frac{1}{2}\log\left(\frac{1+x}{1-x}\right)$

Th Fisher Z-score test is a non-parametric method which does not require prior knowledge of the underlying data distribution and can easily be computed. On the downside, it provides poor results for small sample sizes as well as for detecting small variations in correlation coefficients. It also only provides a one-tail test, limiting its flexibility.

The Jennrich test[2] is a method for testing the equality of two or more correlation derived from independent samples. The test based on the following statistic as summarized by Atiany et al[7]:

With $m$ samples of size $n_1, n_2, \dots, n_m$ drawn from $p$-variate distributions $N_p(\mu_k, \Sigma_k)$ and associated $\Omega_k$ covariance matrices for $k\ in\ [1 \dots m]$. The Jennrich statistic tests for the hypothesis $H_0 : \Omega_1 = \cdots = \Omega_m$ versus $H_1\colon \Omega_i \neq \Omega_j$ for at least one $(i, j)$ is different. The statistic is given as :

$$J = \sum_{i=1}^{m} \left(\frac{1}{2} Tr\left(Z_i^2\right) - (Z_d)^t W^{-1} Z_d\right)$$

where :

i. $Z_i = n_i^{\frac{1}{2}} R_p^{-1}\left(R_i - R_p\right)$
ii. $R_i$ is the correlation of the i-th sample
iii. $R_p$ is the average across of all sample correlation matrices
iv. $W = I + R_p * R_p^{-1}$, with $*$ being the Hadamard product (term by term product of two matrices)
v. $Z_d$ is a diagonal of $Z_i$
vi. $I_p$ is the identity matrix of size $(p \times p)$

The statistic $J$ is asymptotically distributed as a $\chi^2$ with $df = \frac{1}{2}(m-1)p(p-1)$ degrees of freedom.

The Jennrich test is also non parametric and provides an easy $\chi^2$ approximation for high sample size. On the downside it can be computationally heavy and is sensitive to outliers in the data.

All those methods approach the study of difference in correlations between two outcomes through their corresponding correlation matrices. But more recently some papers[8,9] started studying the correlation through the products of two variables, which is the method we present in depth and larger scale in this paper.

##### *Covariance of two outcomes and outcomes product*

Consider a centered predictor $X$, two normally distributed outcomes $Y_1$ and $Y_2$ with mean 0, which depend on an unmeasured variable $U$ capturing the shared variance between $Y_1$ and $Y_2$, and two independent residual terms $\epsilon_1$ and $\epsilon_2$. The covariance between $Y_1$ and $Y_2$ conditional on $X$ is expressed as $Cov(Y_1, Y_2|X) = \mathbb{E}[Y_1 Y_2|X] - \mathbb{E}[Y_1|X]\mathbb{E}[Y_2|X]$. Note that, if not specified otherwise, the product of vectors indicates element-wise product. We are interested in determining simple generative models inducing a non-zero conditional covariance. To assess the theory of this approach we derived $Cov(Y_1, Y_2|X)$ for three models: (1) in the presence of effects of $X$ on the mean of the outcome, (2) in the presence of effects of $X$ on the variance of the outcome, and (3) interaction effects between $X$ and the shared factor $U$ on the mean of the outcome.

In model (1), we assume the predictor $X$ has an effect on the mean of $Y_1$ and $Y_2$:

$$Y_1 = \alpha_1 U + \beta_1 X + \epsilon_1$$

$$Y_2 = \alpha_2 U + \beta_2 X + \epsilon_2$$

The components of the covariance equal:

$$\begin{aligned}
\mathbb{E}[Y_1 Y_2|X = x] &= \mathbb{E}[(\alpha_1 U + \beta_1 X + \epsilon_1)(\alpha_2 U + \beta_2 X + \epsilon_2)|X = x] \\
&= \mathbb{E}[\alpha_1\alpha_2 U^2 + \beta_1\beta_2 X^2 + (\alpha_1\beta_2 + \alpha_2\beta_1)UX + \alpha_1 U\epsilon_1 + \alpha_2 U\epsilon_2 + \beta_1 X\epsilon_2 + \beta_2 X\epsilon_1 + \epsilon_1\epsilon_2|X = x] \\
&= \mathbb{E}[\alpha_1\alpha_2 U^2|X = x] + \mathbb{E}[\beta_1\beta_2 X^2|X = x] + \mathbb{E}[(\alpha_1\beta_2 + \alpha_2\beta_1)UX|X = x] + \mathbb{E}[\alpha_1 U\epsilon_1|X = x] \\
&\qquad + \mathbb{E}[\alpha_2 U\epsilon_2|X = x] + \mathbb{E}[\beta_1 X\epsilon_2|X = x] + \mathbb{E}[\beta_2 X\epsilon_1|X = x] + \mathbb{E}[\epsilon_1\epsilon_2|X = x] \\
&= \alpha_1\alpha_2\mathbb{E}[U^2] + \beta_1\beta_2 x^2 + (\alpha_1\beta_2 + \alpha_2\beta_1)\mathbb{E}[U]x + \alpha_1\mathbb{E}[U]\mathbb{E}[\epsilon_1] + \alpha_2\mathbb{E}[U]\mathbb{E}[\epsilon_2] + \beta_1\mathbb{E}[\epsilon_2]x \\
&\qquad + \beta_2\mathbb{E}[\epsilon_1]x + \mathbb{E}[\epsilon_1]\mathbb{E}[\epsilon_2] \\
&= \alpha_1\alpha_2\mathbb{E}[U^2] + \beta_1\beta_2 x^2,
\end{aligned}$$

$$\begin{aligned}
\mathbb{E}[Y_1|X]\mathbb{E}[Y_2|X] &= \mathbb{E}[\alpha_1 U + \beta_1 X + \epsilon_1|X = x]\mathbb{E}[\alpha_2 U + \beta_2 X + \epsilon_2|X = x] \\
&= (\alpha_1\mathbb{E}[U] + \beta_1 x + \mathbb{E}[\epsilon_1])(\alpha_2\mathbb{E}[U] + \beta_2 x + \mathbb{E}[\epsilon_2]) \\
&= \beta_1\beta_2 x^2,
\end{aligned}$$

It follows that the covariance does not depend on $X$ in this model:

$$\begin{aligned}
Cov(Y_1, Y_2|X = x) &= \alpha_1\alpha_2\mathbb{E}[U^2] \\
&= \alpha_1\alpha_2 var(U)
\end{aligned}$$

In model (2), we assume $X$ has an effect on the variance of one outcome, which we expressed through a latent variable $A$, normally distributed and centered:

$$Y_1 = \alpha_1 U + \beta_1 AX + \epsilon_1$$

$$Y_2 = \alpha_2 U + \epsilon_2$$

The components of the covariance equal:

$$\begin{aligned}
\mathbb{E}[Y_1 Y_2|X = x] &= \mathbb{E}[(\alpha_1 U + \beta_1 AX + \epsilon_1)(\alpha_2 U + \epsilon_2)|X = x] \\
&= \mathbb{E}[\alpha_1\alpha_2 U^2 + \alpha_1 U\epsilon_2 + \alpha_2\beta_1 AXU + \beta_1 AX\epsilon_2 + \alpha_2 U\epsilon_1 + \epsilon_1\epsilon_2|X = x] \\
&= \alpha_1\alpha_2\mathbb{E}[U^2] + \alpha_1\mathbb{E}[U]\mathbb{E}[\epsilon_2] + \alpha_2\beta_1\mathbb{E}[AU]x + \alpha_2\mathbb{E}[U]\mathbb{E}[\epsilon_1] + \mathbb{E}[\epsilon_1]\mathbb{E}[\epsilon_2] \\
&= \alpha_1\alpha_2\mathbb{E}[U^2],
\end{aligned}$$

$$\mathbb{E}[Y_1|X = x]\mathbb{E}[Y_2|X = x] = \mathbb{E}[\alpha_1 U + \beta_1 AX + \epsilon_1|X = x]\mathbb{E}[\alpha_2 U + \epsilon_2|X = x]$$

$$= (\alpha_1 \mathbb{E}[U] + \beta_1 \mathbb{E}[A]x + \mathbb{E}[\epsilon_1])(\alpha_2 \mathbb{E}[U] + \mathbb{E}[\epsilon_2])$$

$$= 0 \,,$$

and,

$Cov(Y_1, Y_2|X) = \alpha_1\alpha_2 var(U)$ which is independent of $X$.

<u>In model (3)</u>, we assume $X$ has an interaction effect with the shared factor $U$ on one outcome:

$$Y_1 = \alpha_1 UX + \epsilon_1$$

$$Y_2 = \alpha_2 U + \epsilon_2$$

It follows that:

$$\mathbb{E}[Y_1Y_2|X = x] = \mathbb{E}[(\alpha_1 UX + \epsilon_1)(\alpha_2 U + \epsilon_2)|X = x]$$

$$= \mathbb{E}[\alpha_1\alpha_2 U^2 X + \alpha_1 UX\epsilon_2 + \alpha_2 U\epsilon_1 + \epsilon_1\epsilon_2|X = x]$$

$$= \alpha_1\alpha_2\mathbb{E}[U^2]x + \alpha_1\mathbb{E}[U]\mathbb{E}[\epsilon_2]x + \alpha_2\mathbb{E}[U]\mathbb{E}[\epsilon_1]x + \mathbb{E}[\epsilon_1]\mathbb{E}[\epsilon_2]$$

$$= \alpha_1\alpha_2\mathbb{E}[U^2]x,$$

$$\mathbb{E}[Y_1|X = x]\mathbb{E}[Y_2|X = x] = \mathbb{E}[\alpha_1 UX + \epsilon_1|X = x]\mathbb{E}[\alpha_2 U + \epsilon_2|X = x]$$

$$= (\alpha_1\mathbb{E}[U]x + \mathbb{E}[\epsilon_1])(\alpha_2\mathbb{E}[U] + \mathbb{E}[\epsilon_2])$$

$$= 0$$

and,

$Cov(Y_1, Y_2|X = x) = \alpha_1\alpha_2 x \; var(U)$

In this model we can see a direct effect of the predictor X on the value of the covariance.

<u>Note on model 1:</u>

Our product-based approach corresponds to the evaluation of the first term of the covariance estimator, *i.e.* $\mathbb{E}[Y_1Y_2|X = x]$. In model 2 and 3, the product of the conditional expectation ($\mathbb{E}[Y_1|X = x]\mathbb{E}[Y_2|X = x]$), and our product-based approach corresponds exactly to the covariance between $Y_1$ and $Y_2$. However, in model 1, this second term is not null, and our product-based approach carries the additional term $\beta_1\beta_2 x^2$.

Let's denote $P$ the vector of element-wise product of $Y_1$ and $Y_2$. From model (1) description, the estimated regression coefficient of $P$ on $X$, denoted $\hat{\gamma}$, and obtained from the standard linear model: $P \sim \gamma_0 + \gamma X$ equals:

$$\mathbb{E}[\hat{\gamma}] = \mathbb{E}\big[\mathbb{E}[\hat{\gamma}|X = x]\big]$$

$$= \mathbb{E}\big[\mathbb{E}[PX|X = x]\big]$$

$$= \mathbb{E}[\alpha_1\alpha_2 x\mathbb{E}[U^2] + \beta_1\beta_2 x^3]$$

$$= \beta_1\beta_2\mathbb{E}[X^3]$$

As the element-wise product of two centered normal is centered, and the third moment of a normal is also centered, when all variables are normally distributed and centered, one can notice that the independence between all variables implies that $\mathbb{E}[\hat{y}] = 0$.

However, when the predictor is not normally distributed, the expected value of $\hat{y}$ becomes non-trivial. For example, assume $U$, $\epsilon_1$and $\epsilon_2$ are normally distributed, but $X$ is a genetic variant coded additively, and thus following a binomial distribution $X$~$\mathcal{B}(n,p)$. All terms will again be equal to 0, except for $\mathbb{E}[\boldsymbol{X}^3]$, the third moment of $X$, which equals $n(n-1)(n-2)p^3 + 3n(n-1)p^2 + np$. Importantly, note that this difference between the estimated effect based on the product-terms and the true effect on the covariance, requires effect of the predictor on both outcomes (i.e. $\beta_1 \neq 0$ and $\beta_2 \neq 0$), and would be negligible unless both $\beta_1$ and $\beta_2$ are substantially large.

<u>Note on model 2:</u>

Our product-based approach assesses differences in the covariances between two outcomes $Y_1$ and $Y_2$. When $Y_1$ and $Y_2$ are standardized, effect on the covariance can in general be transpose to effect on the correlation. However, in the special case of model 2, where the predictor is associated with the variance of $Y_1$, this equality is not valid anymore, as the correlation depends on $X$, while the covariance does not:

$$Cor(Y_1, Y_2|X = x) = \frac{Cov(Y_1, Y_2|X = x)}{\sqrt{var(Y_1|X)var(Y_2|X = x)}}$$

$$= \frac{\alpha_1\alpha_2 var(U)}{\sqrt{var(Y_1|X = x)}}$$

$$= \frac{\alpha_1\alpha_2 var(U)}{\sqrt{var(\alpha_1 U + \beta_1 AX + \epsilon_1|X = x)}}$$

$$= \frac{\alpha_1\alpha_2 var(U)}{\sqrt{\alpha_1^2 var(U) + var(\epsilon_1) + \beta_1^2 var(A)x}}$$

Note that the scaling of the outcomes does not impact this difference.

##### ***Miscalibration of the sum of PCs test***

MANOCCA is readily applicable to the matrix of outcome product in all scenarios where $p$ is substantially smaller than $N$. However, in many omics' datasets, the number of products, which is quadratic with the number of outcomes $k$, can be substantially larger than the sample size and a multivariate test of all the products would not be applicable. We used Principal Components Analysis (PCA) to reduce the dimension of $P$, using the top principal components as the primary outcome. Because principal components (PCs) are independent from each other by construction, the chi-squares for association obtained from the top $m$ principal components could, in theory, be summed to form a joint statistic $T = \sum_m \chi_i^2$ that follows a chi-square distribution with $m$ degrees of freedom. However, this test is strongly miscalibrated when the number of PCs analysed jointly is not of several orders of magnitude smaller than the actual sample size. This bias is due to a deviation of the variance of $T$ from the expected under the null (**Fig. S1a-c**) and is driven by a realized covariance between PCs smaller than expected under the null (**Fig. S2**). To solve this issue, we implemented a Wilks test of PCs that addresses the miscalibration of the sum of chi-squared statistics (**Fig. S1d**).

#### *Extended description of the Milieu Intérieur data used for the screenings*

The *Milieu Intérieur* (MI) Consortium is a population-based cohort initiated in September, 2012[10]. It comprises 1,000 healthy volunteers from western France, with a 1:1 sex ratio. The cohort collected a broad range of variables, including genomic, immunological, environmental, and clinical outcomes. We conducted systematic MANOCCA screenings for environmental effects on the covariance of two sets of data: 169 flow cytometry-based immune cell phenotypes and 33 health-related blood biomarkers, including 22 metabolites and 11 cell counts. We focused on two types of predictors: health and lifestyle factors collected from questionnaires, and genome-wide variants. Except when used as predictor, all analyses were adjusted for age, sex and body mass index (BMI). For blood metabolites, the number of products allowed for a direct analysis of the products without requiring the PCA step, and we considered both the products and the PCs as outcomes. For comparison purposes, we also conducted, for each screening, a standard MANOVA on the mean of the multivariate outcome.

Immune cell phenotypes were derived for 1,000 participants from whole blood using ten flow cytometry panels capturing major cell populations present in human blood: granulocytes (neutrophils, basophils and eosinophils), monocytes, natural killer (NK) cells, dendritic cells (DCs), innate lymphoid cells (ILCs), T cells (γδ T cells, mucosa-associated invariant T cells, NKT cells, $T_{reg}$ cells and helper T cells) and B cells[11]. The 169 immunophenotypes included 79 absolute counts of circulating cells, 87 expression levels of cell-surface markers (quantified as mean fluorescence intensity (MFI)), and 3 ratios of cell counts or MFI values (**Table S1**). The 22 blood metabolites and 11 blood cell counts were measured by standardized clinical lab tests[10] (**Table S2**).

*Milieu Intérieur* volunteers were asked to fill in a questionnaire covering multiple panels. Among those available, we pre-selected a set of 122 numeric variables related to demographics, medical history, vaccination history, psychological traits, nutrition, socio-professional information, smoking habits and physiological measurements. This included the top three diet factors derived from the Nutrinet[12] study. We filtered variables with over 50% missing values, and binary predictors with a frequency smaller than 10%, leading to a total of 49 factors available and a sample size up to 992 individuals. Sample sizes available per variable tested and descriptive statistics are provided in **Table S3**.

Finally, we also conducted a genome-wide genetic screening for the same two sets of outcomes to demonstrate the scalability of our approach. Blood sample was collected for all participants and genotypes were derived using an *Illumina HumanOmniExpress-24 array* for all 1,000 participants. Imputation was conducted using IMPUTE2[13] using the Haplotype Reference Consortium (release 1.1) as a reference panel[14] for 956 participants. After removing variants with an imputation info score < 0.8, and minor allele frequency below 5%, a total of 5,667,803 genotyped and imputed variants remains for analyses. We ran a principal component analysis on the genotype matrix using Plink2[15] and removed 62 ancestral outliers individual (i.e. individuals whose genetic makeup diverges from the rest of the population studied based on a principal component plot[16,17]), resulting in a total sample size of up to 894 participants from whom genetic and phenotypic data were available.

#### Supplementary Figures

##### *Figure S1: Miscalibration of a joint test of PCs based on the sum of chi-squared*

We assessed the calibration of the joint test of multiple PCs based on the sum of individual chi-squared as a function of the number of PCs kept. We first simulated series of replicates with a sample size $N$ in [350; 1,000; 2,000; 10,000], each including a multivariate normal outcome $\boldsymbol{Y}$ of size $N \times 100$ and a predictor $X \sim B(0.4)$. For each replicate, we computed the product matrix $\boldsymbol{P}$ from $\boldsymbol{Y}$, applied a principal component analysis on $\boldsymbol{P}$, and derived a one degree of freedom chi-square $\chi^2_{1\mathrm{df},i}$ for association between $X$ and each of the top $m$ principal component $PC_{i=1\ldots m}$ from a standard linear regression. We formed joint statistics $S$ from the top PCs test as the sum: $S = \sum_{i=1\ldots m} \chi^2_{1\mathrm{df},i}$, which is expected to follows a $m$ degree of freedom chi-squared under the null. Panel a) and b) present the mean and variance of $S$, respectively, against their expected value, illustrating the correct calibration of the mean and the miscalibration of the variance of $S$. Panel c) presents the *P*-value distribution of the sum of chi-square test for the scenario including 300 PCs and $N = 1000$. The Kolmogorov–Smirnov test for deviation from a uniform [0,1] distribution, was highly significant in those data (*P* < 1x10$^{-16}$). Panel d) presents the *P*-value obtained from the same data when applying a Wilks test on the same PCs. The Kolmogorov–Smirnov test for deviation from a uniform [0,1] distribution was not significant (*P*=0.94).

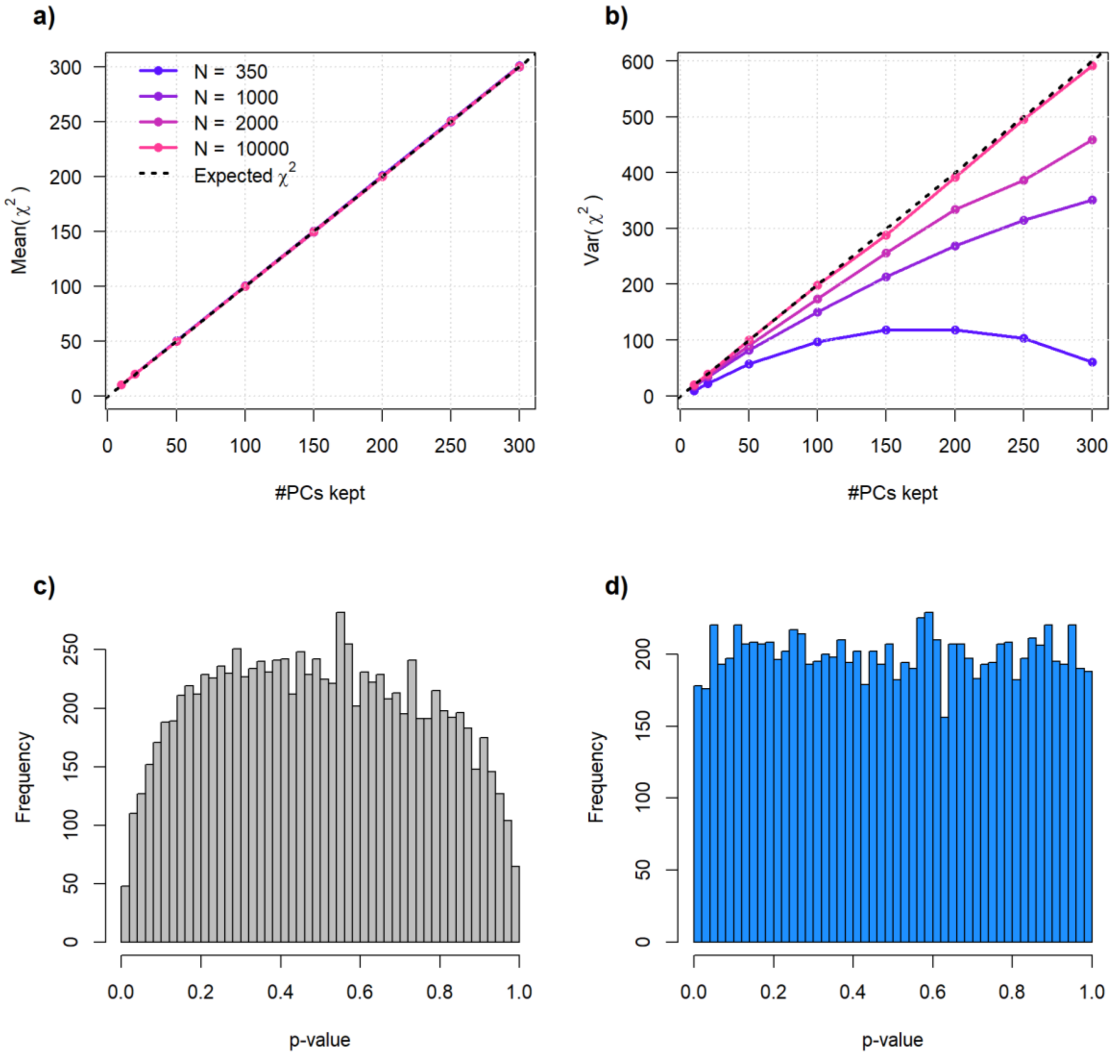

***Figure S2: Relationship between sample size and bias in the joint test of PCs***

We used a simple two variables model to illustrate the cause of the bias for the joint test of principal component based on the sum of chi-squared. We simulated series of replicates, each including two independent normally distributed variables $Y_1$ and $Y_2$ and 1,000 predictors for a sample size varying in [5; 100]. For each replicate we derived the chi-squared for association between the two outcomes and the predictors, $\chi_1$ and $\chi_2$. For each sample size, we derived the covariance between the resulting chi-squared (a) and the two intermediate terms, *i.e.* the expected value of their product $E[\chi_1\chi_2]$ (b), and the product of the expected value $E[\chi_1]$ $E[\chi_2]$ (c). We plotted the same parameters but using the principal components $PC_1$ and $PC_2$ derived from $Y_1$ and $Y_2$. While the expected chi-squared of each PCs are well calibrated, and follow the same distribution as the original independent variables (c), the expected value of the product of chi-square from the PCs is substantially smaller than expected by chance (b) for small sample size. It explains the negative bias for the covariance observed for PCs, not observed for the original outcomes (panel a).

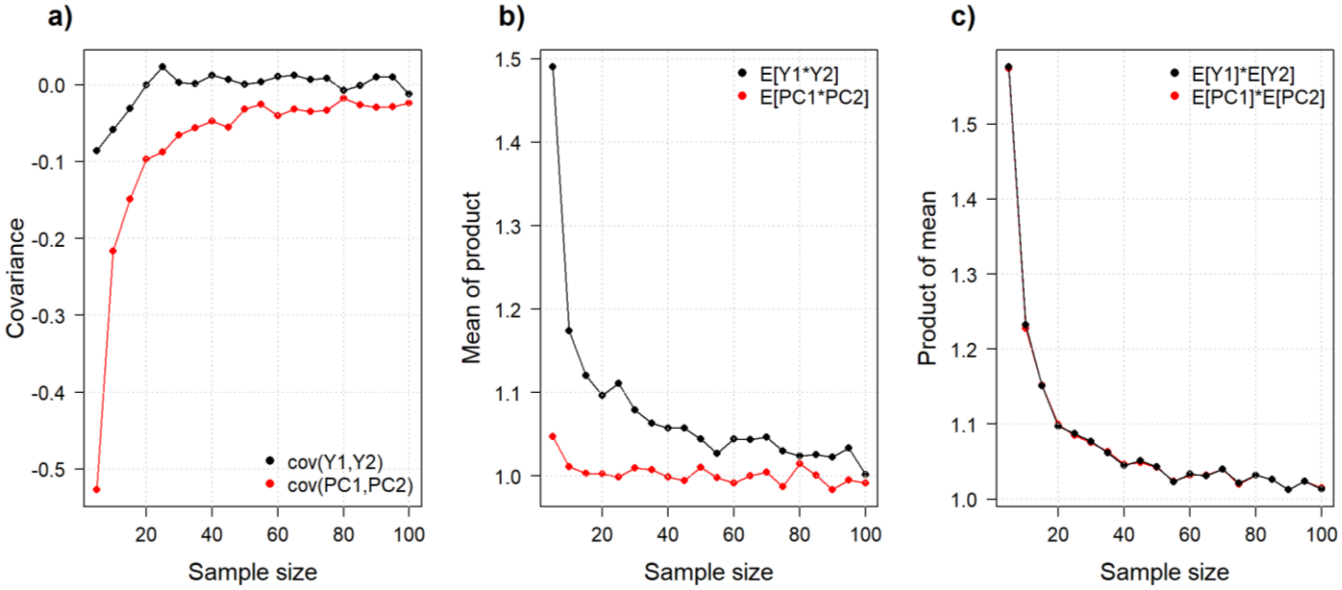

***Figure S3. Calibration for binary and continuous predictors across pre-processing and PC***

We simulated series of 100 replicates, each including 1,000 individuals and a multivariate outcome $\mathbf{Y}$ mimicking OMICs data. The outcome $\mathbf{Y}$ included 400 variables drawn from multivariate chi-squared. For each replicate we draw 1,000 predictors and performed the proposed MANOCCA test while applying no transformation, a rank inverse normal transformation on the product matrix, a rank inverse normal transformation on the principal component matrix, or a rank inverse normal transformation on both. The validity of the test was assessed using a Kolmogorov–Smirnov test for deviation from a uniform [0,1] distribution of the P-value across the 1,000 predictors tested, while varying the number of PCs used. Panel a) present the results when using a normally distributed predictor. Panel b) present the results when using a binary outcome with a frequency of 0.05.

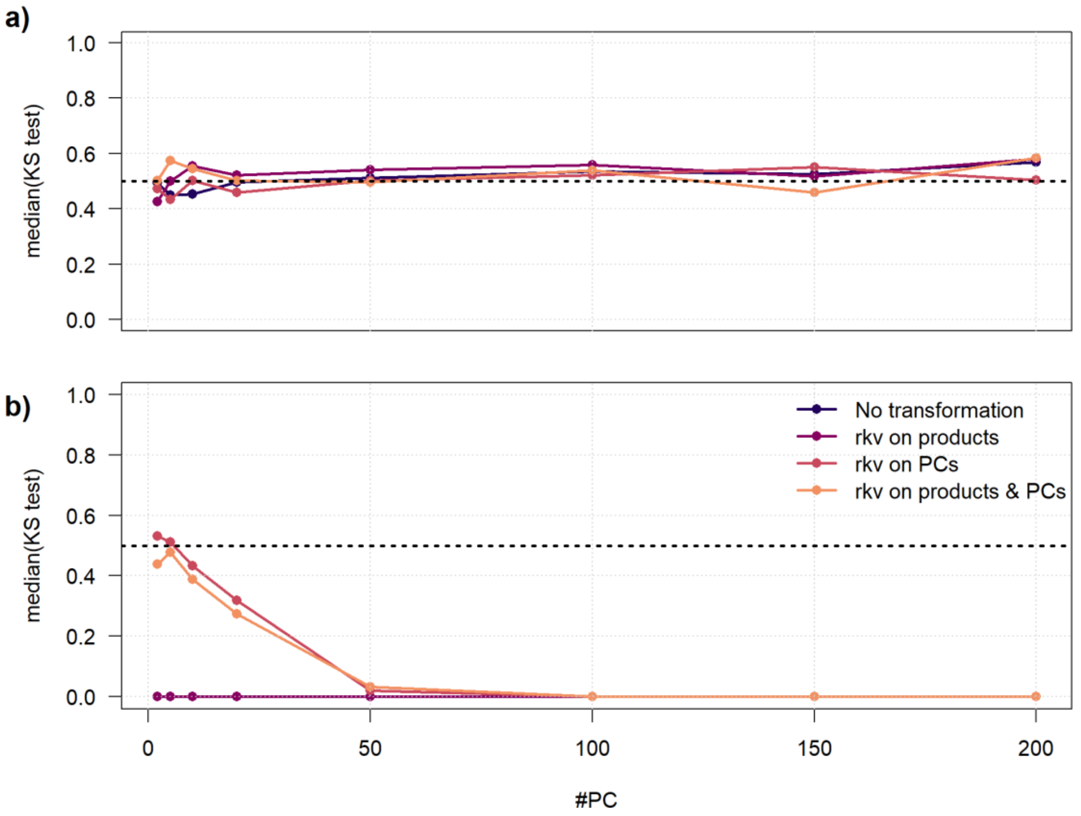

***Figure S4. Test calibration as a function of the number of PCs tested***

We simulated series of 1,000 replicates where we generated data of size $N$ and k variables from a covariance matrix derived from 169 real blood biomarkers. For each simulation, 1,000 random binary predictors, and in one case genetic predictor, with frequency f in [0.01, 0.05, 0.1, 0.4] were generated. Finally, we derived a MANOCCA p-value for each predictor on a range of PC kept [1,…10, 20, … 100, 200] and used a Kolmogorov-Smirnov uniformity test to assess the p-values distribution and therefore the calibration of the test. Panel a) displays the standard screening scenario using N = 1000, k = 169, b) shows N = 1000, k = 30, c) shows N = 1000, k = 169 but with a genotype predictor (0,1,2), d) shows N = 5000, k = 169.

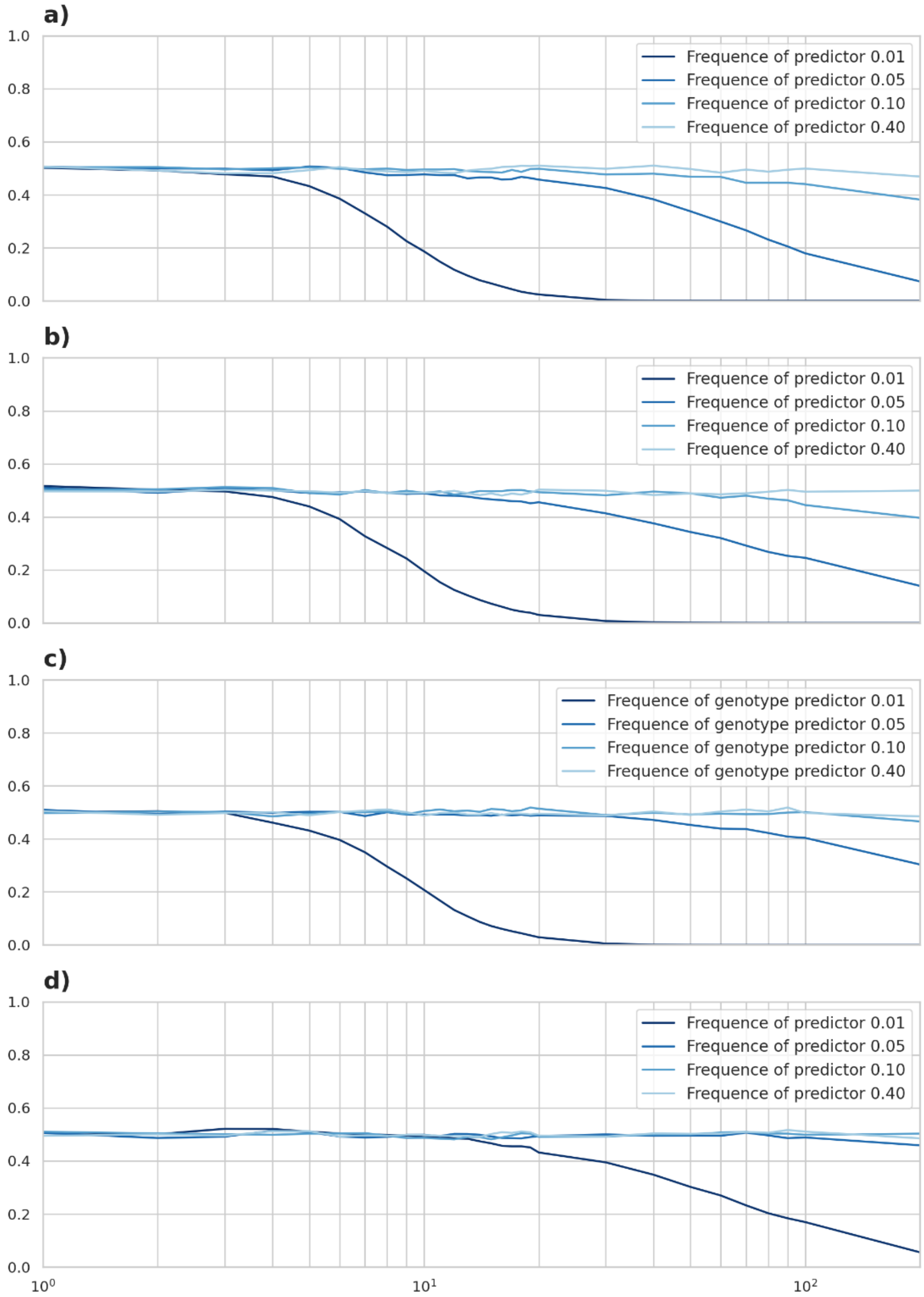

***Figure S5. Guideline for Principal Component selection***

Data was generated using 1000 replicates of varying sample size N [1000 - 5000] and binary predictor $X$ frequencies [0.01-0.09, 0.1-0.5]. For each pair (N, frequency, $Pvalue_{KS\ test}$) we selected the last number of PC where the mean P-value for the Kolmogorov-Smirnov test over the 1000 replicates was above the threshold of 0.47. The simulations were ran until 400 PCs were kept leaving a saturation threshold at 400 PC kept for the MANOCCA test. The x-axis displays the ratio frequency*sample size, while the y-axis displays the number of PC kept.

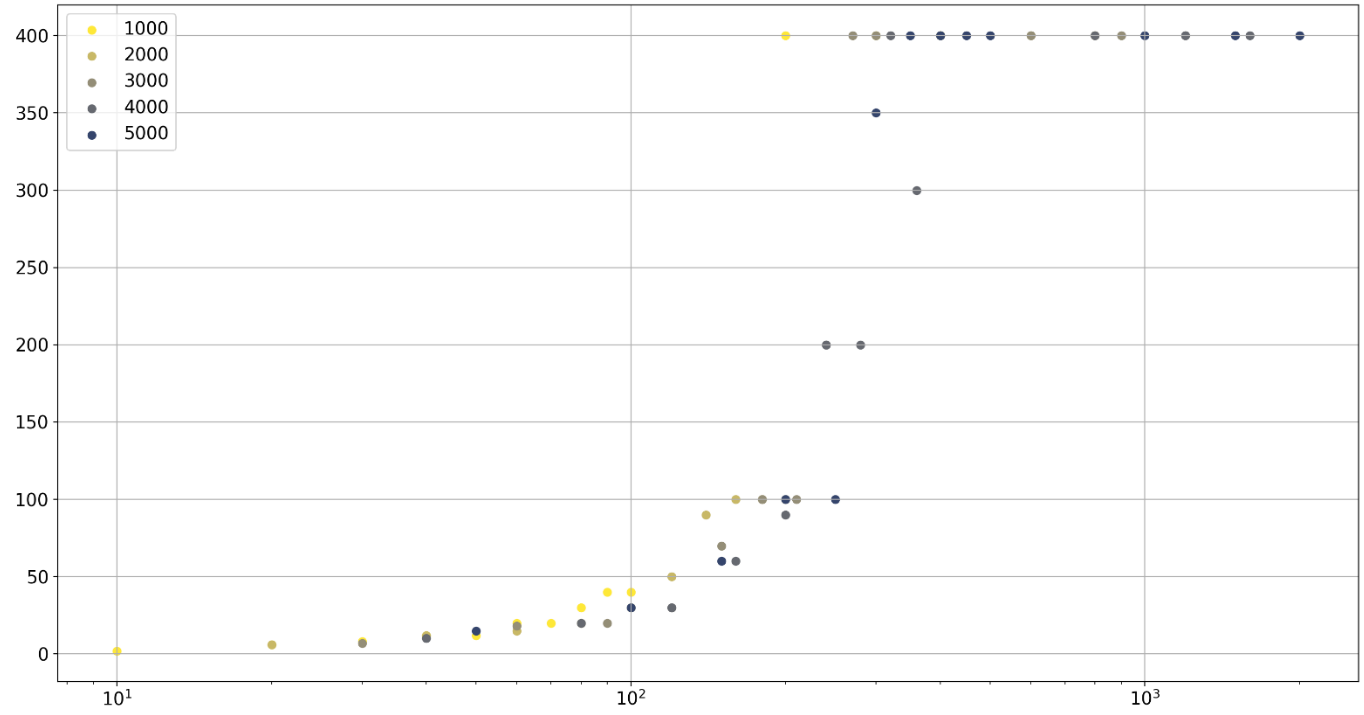

***Figure S6. Correlation matrices in Milieu Interieur***

Correlation matrices between 169 flow cytometry immunity phenotypes (a) and between 33 blood biomarkers (b) derived from 992 healthy individuals from the Milieu Intérieur. Columns and lines were ordered using the default *hclust* function from the stat package in R. Panels c) and d) present the distribution of the correlation for the flow cytometry and blood biomarkers, respectively.

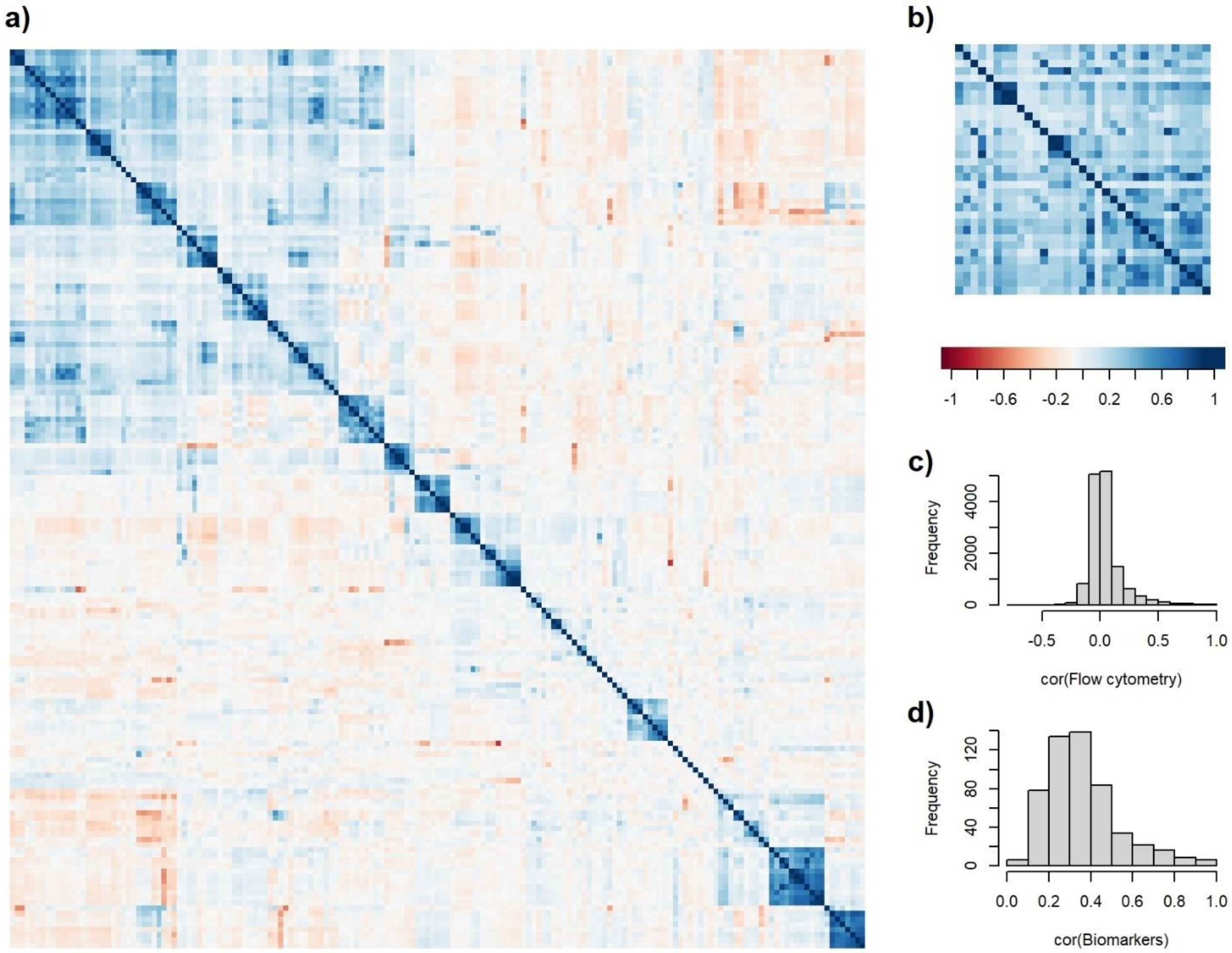

***Figure S7. Correlation between PCA-based and product-based MANOCCA***

We screened for the effect of 49 health and lifestyle factors on the covariance of 33 blood biomarkers using the MANOCCA approach. We applied MANOCCA directly to the products matrix of the biomarkers, and to the top principal components (PCs) of the product matrix. The figure presents the correlation between P-value obtained from the MANOCCA applied to the products and the PCs while varying the number of PCs from 5 to 500 with a step of five.

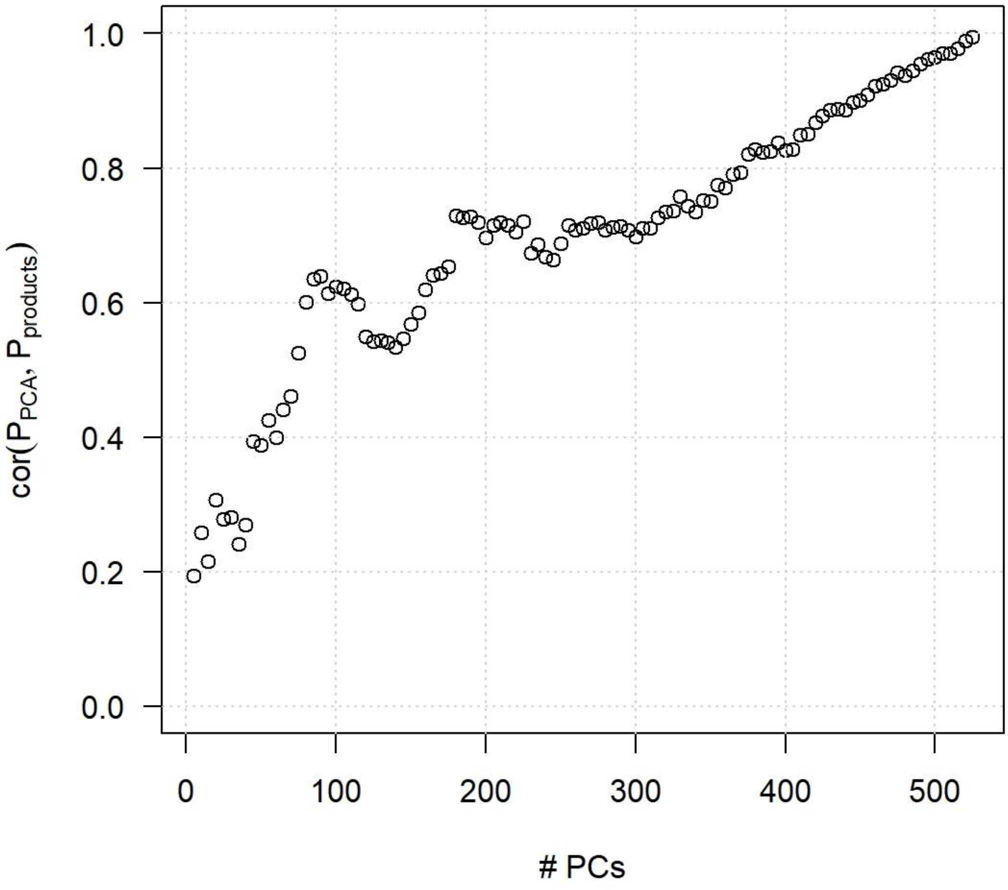

***Figure S8. Correlation between MANOCCA and Manova association***

We compared the association signal obtained from our MANOCCA screenings using the 49 health and lifestyle factors with the results from a MANOVA screening on the same outcomes and predictors sets. Comparison was conducted using the -log10(*P*-value) of Manova, and the -log10(*P*-value) of the minimum *P*-value from MANOCCA over the 40 sets of principal components considered (from 5 to 200 with a step of 5). Panel a) displays the results for the analysis of the 169 flow cytometry immunity phenotypes, and panel b) displays the results for the analysis of the 33 blood biomarkers.

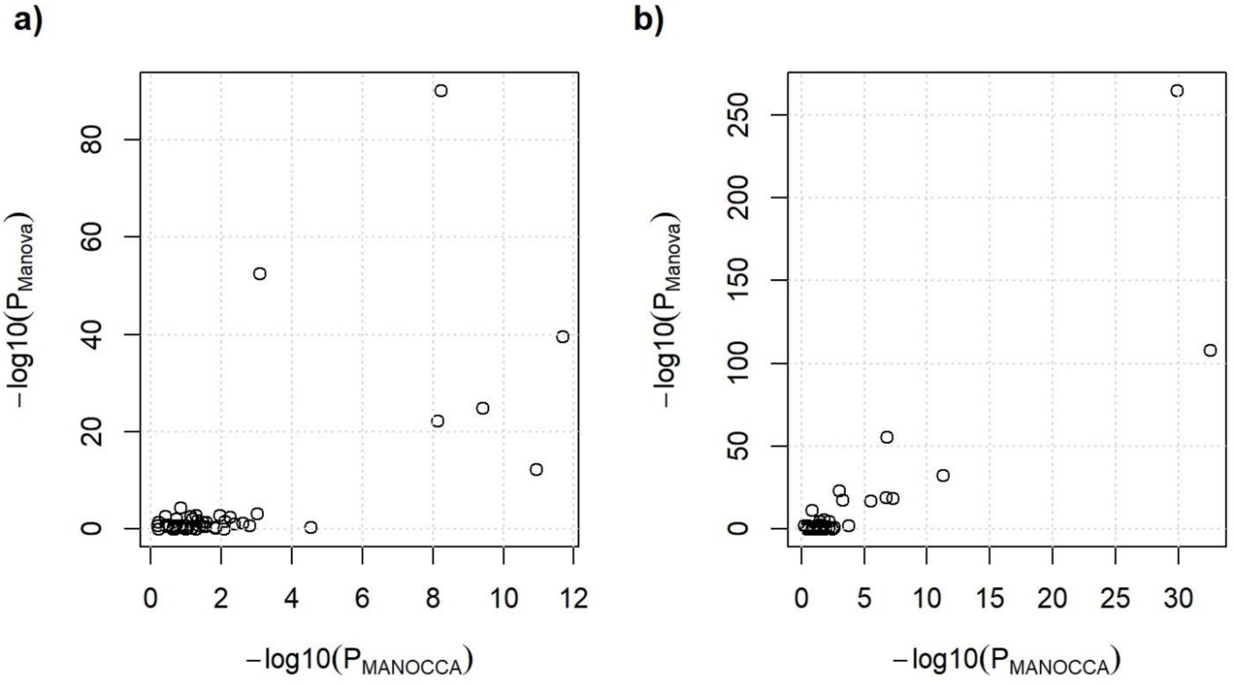

*Figure S9. Blood biomarkers and flow cytometry GWAS*

We ran genome-wide association studies (GWAS) using MANOCCA on 5,699,237 variants with a MAF > 5% in 890 samples with complete genetic and phenotypic data. Panel a) display results for the screening of the covariance of the 33 blood biomarkers. Panel b) display the results for the screening of the covariance of the 169 flow cytometry immunity phenotypes. We varied the number of principal components (PCs) from 5 to 50 with a step of 5 resulting in for a total of 10 GWAS per dataset. Left panels display the Manhattan plot of the minimum *P*-value per variant over the 10 GWAS. The red dash line indicates the stringent Bonferroni correction threshold of 5 x $10^{-9}$ accounting for the 40 sets of principal components considered. Right panels display the QQplot for each of the 20 GWAS. The mean lambda value across the 20 GWAS equal 0.996 and 1.001 for the blood biomarkers and flow cytometry data, respectively.

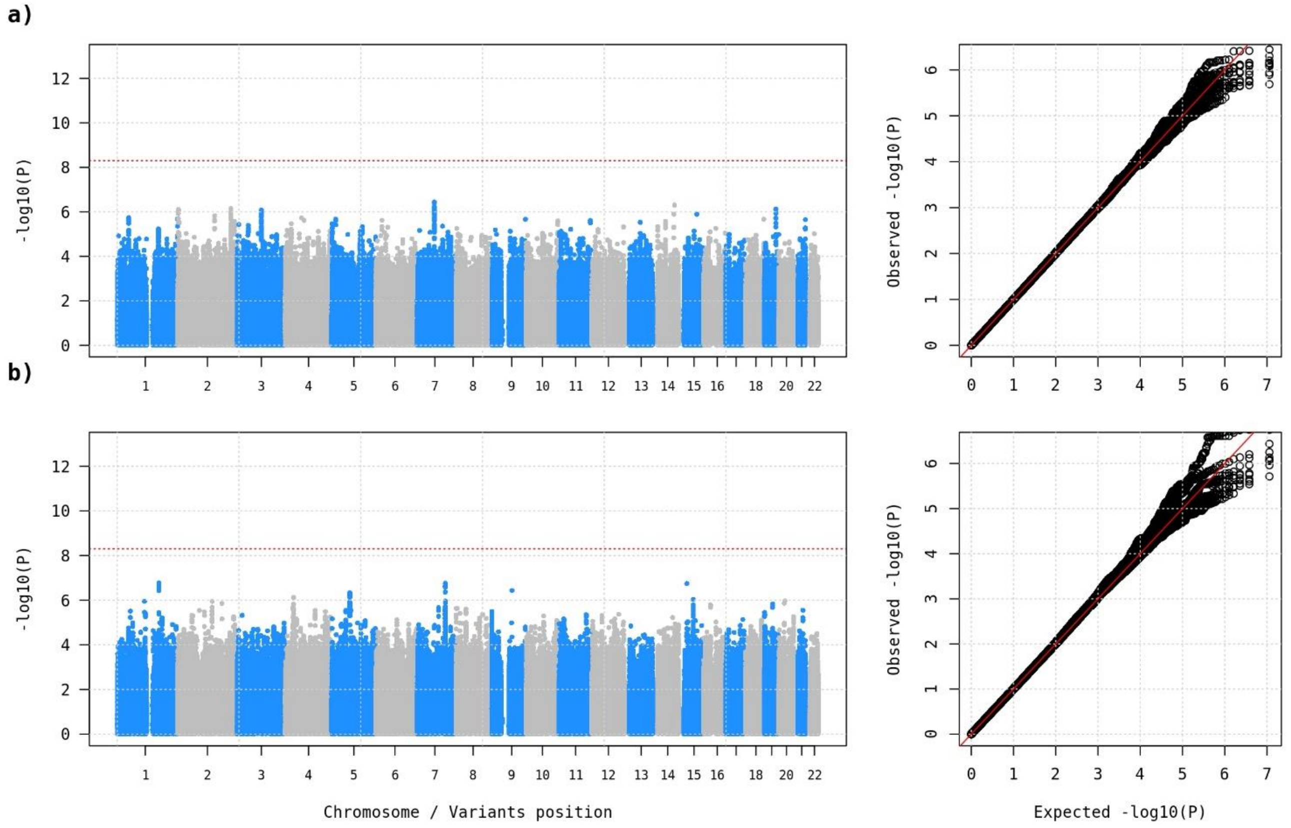
